# Aromatic amino acids in the orthosteric region regulate cannabinoid receptor 1 conformation transitions

**DOI:** 10.1101/2025.08.18.670779

**Authors:** Cecylia S. Lupala, Xuanxuan Li, Xuefei Li, Haiguang Liu

## Abstract

Cannabinoid receptor 1 (CB1) is a class A G protein–coupled receptor that exhibits constitutive activity by spontaneously sampling both active and inactive states in the absence of ligand binding. Structural studies have identified a domain of aromatic residues, F200^3.36,^ W356^6.48^, and W279^5.43^, that modulates receptor function and ligand engagement; however, their mechanistic roles remain unclear. Here, we use equilibrium and targeted molecular dynamics simulations to investigate how alanine substitutions at these positions alter CB1 dynamics. Our results show that mutations, especially the F200A+W356A combination, enhanced conformational flexibility and shifted the receptor toward inactive-like states. W356^6.48^ emerged as a structural pivot essential for active-state stability, F200^3.36^ acted as a late-stage kinetic barrier to inactivation, and W279^5.43^ modulated orthosteric pocket packing and flexibility, amplifying destabilization when combined with F200^3.36^ or W356^6.48^ mutations. Targeted simulations showed that mutation of this aromatic domain lowers energetic barriers and rewires the inactivation pathway. These findings define the structural logic of CB1 toggle-switch control and provide a mechanistic framework for designing modulators that tune basal activity in CB1.

## Introduction

G protein-coupled receptors (GPCRs) constitute the largest and most diverse family of membrane proteins in the human genome and represent over one-third of current drug targets^1–3^. These receptors mediate signal transduction through conformational transitions between inactive and active states triggered by ligand binding. While the canonical activation mechanism involves outward displacement of transmembrane helix 6 (TM6) and intracellular rearrangements enabling G-protein engagement^4–6^, emerging studies have revealed noncanonical features and state-dependent flexibility across GPCR subclasses^7–10^. CB1 receptor is a class A GPCR highly expressed in the central nervous system, regulates diverse physiological processes including synaptic plasticity, pain sensation, and energy homeostasis^11,12^. CB1 responds to endogenous cannabinoids, phytocannabinoids like Δ9-tetrahydrocannabinol (THC), and synthetic ligands, making it a prominent therapeutic target for disorders such as obesity, addiction, and neuropathic pain^13–15^.

Unlike many class A GPCRs, CB1 displays constitutive activity and adopts multiple conformational states even in the absence of ligands^16–19^. Experiments have shown that CB1 receptors exist in both inactive (RG_GDP_, G-protein pre-coupled) and active (RG_GTP_) states^18^. Generally, an agonist or partial agonist alters the energy landscape of CB1 conformation space, favoring the conformation of the active state, and effectively shifting the structures towards active states upon agonist binding. Likewise, inverse agonists shift the conformations towards the inactive state when bound to the receptor. Apart from ligand or G-protein binding, mutations in the residues involved in GPCR activation can also modify the equilibrium toward either the active or inactive states. For example, constitutively active mutations increase the basal activity of the receptor in the absence of a ligand^20–22^. Moreover, experimental studies have shown that the DRY motif, highly conserved among class A GPCRs, plays a central role in CB1 by mediating G-protein activation and β-arrestin engagement through the stabilization of distinct biased receptor conformations. Single alanine substitutions partially impair G-protein signaling, whereas double or triple mutations severely disrupt it, shifting the signaling bias toward β-arrestin pathways^23^.

Structural studies have provided insights into the activation mechanisms of CB1, revealing intricate conformational changes that underline receptor signaling^17,24,25^. Crystal structures of CB1 in both active and inactive states reveal unconventional features of receptor activation. Notably, substantial structural rearrangements occur in TM1, TM2, and TM6, consistent with a ‘seesaw’ mechanism characterized by inward movement of the extracellular regions of TM1 and TM2, and outward displacement of the intracellular portion of TM6 in the active conformation relative to the inactive state (**Fig. 1a**). As a result, the activation of CB1 is accompanied by conformational changes that shrink the volume of the orthosteric binding site by 53%, meanwhile enlarging the binding pocket on the cytoplasmic side^16^ (**Fig. 1b**). Notably, CB1 lacks the conserved P-I-F motif and instead employs an alternative mechanism involving two residues F200^3.36^ and W356^6.48^ deep within the orthosteric site^26–28^.

**Fig. 1.**
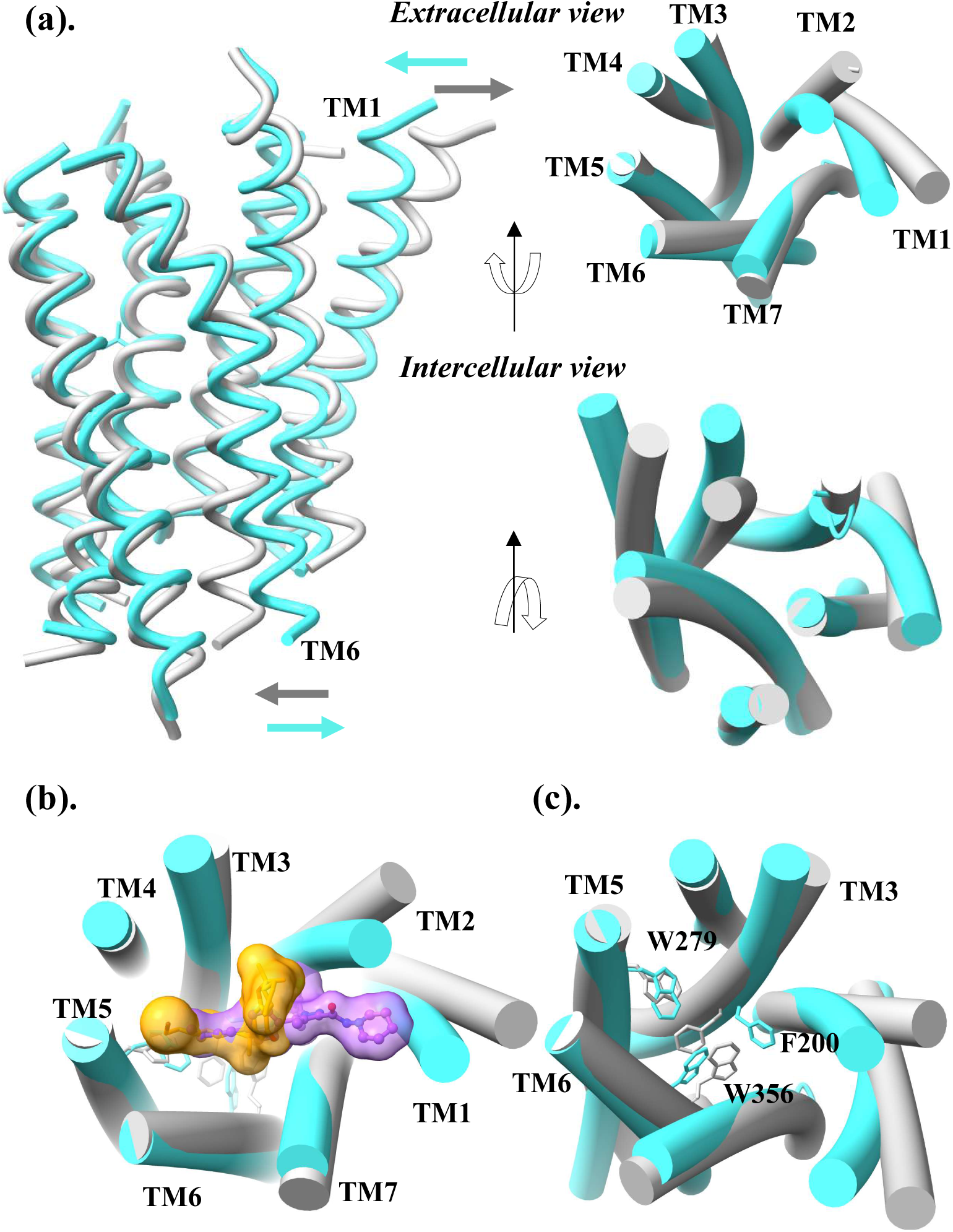
Structural comparison of inactive and active CB1 receptor conformations. **(a)** Superimposed inactive (gray) and active (cyan) CB1 structures highlight key conformational changes upon activation. TM6 shifts outward on the intracellular side in the active state, opening the G-protein binding site, while swinging inward in the inactive state to occlude it. **(b)** On the extracellular side, activation induces inward movements of TM1 and TM2 toward the ligand-binding cavity, reducing the volume of the orthosteric site (shown with a smaller ligand, yellow). **(c)** In the aromatic microdomain, the toggle switch is defined by π-stacking between F200 (TM3) and W356 (TM6) in the active state. W279 (TM5) engages with both residues, contributing to local conformational coupling and stabilization.

In the inactive conformation of CB1, residues F200^3.36^ and W356^6.48^ engage in a π–π stacking interaction (**Fig. 1c**). Together, they form a “twin toggle switch” that plays a central role in coupling orthosteric ligand binding to intracellular signaling via conformational rearrangements. Upon receptor activation, F200^3.36^undergoes a rotameric flip toward TM1, vacating space that permits W356^6.48^ to shift and drive the outward displacement of TM6, a hallmark of GPCR activation^16^. A nearby residue, W279^5.43^, contributes to this aromatic domain, forming interactions with both F200^3.36^ and W356^6.48^ in the inactive state; however, upon activation, W279^5.43^ maintains contact primarily with W356^6.48^. Experimental studies underscore the functional importance of this aromatic domain. The F200A mutation enhances ligand-independent (constitutive) activity, while W356A increases agonist-induced activation^20^. Conversely, W279A has been shown to reduce binding affinity for the inverse agonist Rimonabant by approximately 1000-fold^20,29^. These findings suggest that the twin toggle switch formed by F200^3.36^ and W356^6.48^ is a critical regulator of the CB1 conformational landscape and that mutations within this aromatic cluster can markedly alter the energetic barriers governing activation transitions.

Molecular dynamics (MD) simulations have proven indispensable for studying GPCR activation mechanisms beyond the constraints of crystallography structures, capturing the continuum of conformational states and quantifying transition dynamics^30–35^. Simulations of β2-adrenergic and adenosine A2A receptors have linked rotameric (W^6.48^) toggle switch movements to helical rearrangements during receptor activation^36,37^. For CB1, molecular dynamics simulations investigating the role of the aromatic domain, particularly the toggle switch, have included a bound ligand^38^, while ligand-free studies have generally not incorporated their mutations^39^. As a result, the structural contributions of the aromatic domain remain incompletely understood, especially in the ligand-free state.

In this study, we used equilibrium and targeted all-atom MD simulations to explore CB1 receptor conformational dynamics and the impact of mutations in the aromatic domain in the orthosteric binding site. Equilibrium MD simulations were used to capture fluctuations and assess the stability of the active state, revealing insights into conformational flexibility. Concurrently, targeted MD simulations accelerated transitions to the inactive conformation, enabling evaluation of inactivation propensity and transition pathways. These complementary approaches collectively elucidated the structural plasticity and the critical role of the aromatic cluster in CB1 gating. The aromatic domain residues F200^3.36^, W279^5.43^ and W356^6.48^ were subjected to alanine mutation studies individually, and in all possible combinations. To rule out the ligand binding effects, the resulting 8 systems were simulated in their *apo* form, without agonist or antagonist. As a receptor with tendency to constitutively activate, CB1 receptor provides an opportunity to study the conformation flexibility in the absence of ligands. Structural features derived from CB1 crystal structures were used to quantify conformational changes between the active and inactive states. MD simulations revealed that both wild-type (WT) CB1 and the F200A single mutant largely retained active-like conformations throughout the simulation trajectories. In contrast, the W279A and W356A single mutants exhibited reduced stability of the active state, reflected by intracellular TM6 displacement and extracellular rearrangements of TM1–TM2. Double and triple alanine mutants displayed a marked shift toward inactive-like conformations, indicating a cumulative destabilizing effect. Moreover, targeted MD simulations demonstrated that although the global TM bundle transitions readily, the rotameric switch of F200^3.36^ required elevated force to adopt its inactive conformation, indicating its role as a late-stage conformational barrier. In contrast, disruption of the aromatic domain via alanine substitutions significantly lowered the energetic barriers for full inactivation. Collectively, these findings support a model in which the toggle switch residue F200^3.36^ and the surrounding aromatic domain function as a steric and energetic gate that regulates CB1 activation. Disruption of this gate enhances conformational flexibility and sampling of inactive-like states, providing mechanistic insight into the constitutive activity of CB1 and offering structural principles for designing biased or allosteric GPCR modulators.

## Results

### Mutations in the aromatic domain shift CB1 toward inactivation

To evaluate conformational states, TM backbone RMSD values were calculated after aligning simulated structures to active or inactive CB1 crystal structures. Distributions aggregated from three independent trajectories per system (**Fig. 2**) revealed that WT CB1 remained predominantly active-like, with an average RMSD of ∼2.6 Å to the inactive state, similar to the difference between active and inactive crystal structures (2.75 Å). RMSD to the active state remained ≤0.66 Å (**Fig. S1a**).

**Fig. 2:**
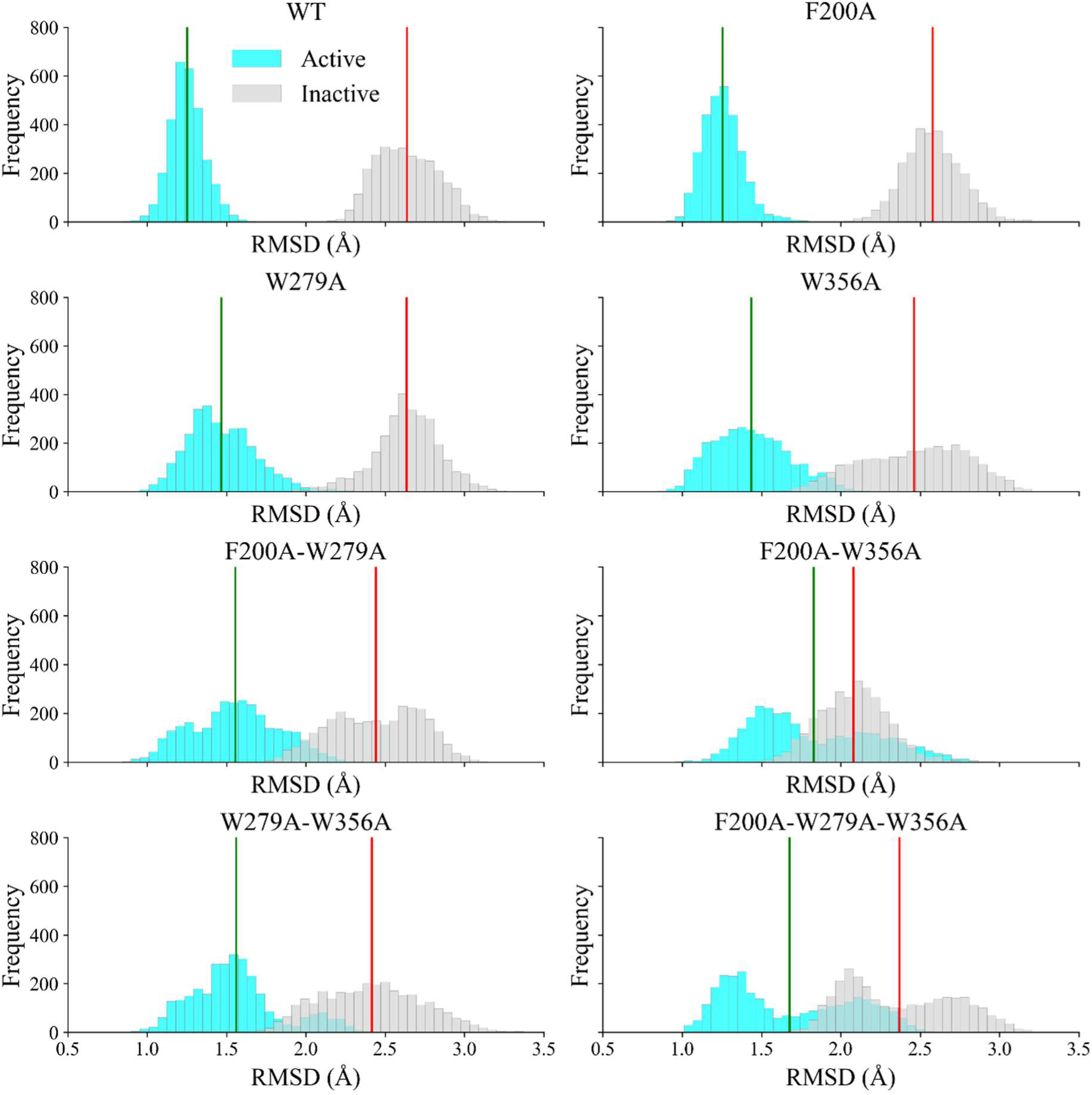
Global conformational shifts relative to active and inactive CB1 states. Backbone RMSD distributions of the transmembrane (TM) region from 1000 ns molecular dynamics simulations of wild-type (WT) and mutant CB1 receptor systems, relative to experimental active (cyan) and inactive (gray) structures. Vertical lines indicate mean RMSD values: green for the active reference and red for the inactive reference. WT and single mutants (F200A, W279A) predominantly align with the active conformation, while double and triple mutants show a progressive shift toward the inactive state. Notably, the F200A+W356A mutant displays the lowest mean RMSD relative to the inactive structure, indicating a faster and more stable transition to inactivation.

Using WT as a reference, the W356A single mutant, all three double mutants (F200A+W279A, F200A+W356A, W279A+W356A), and the triple mutant shifted toward the inactive state, showing reduced RMSD to the inactive conformation. These substitutions replace bulky aromatics with smaller alanines, weakening orthosteric interactions, increasing flexibility, and promoting sampling away from the active state. The F200A+W356A mutant transitioned fastest, with the lowest average RMSD (1.9 Å) and closest match to inactive CB1 (1.39 Å). F200A and W279A alone showed no strong inactivation tendency, and their active-like states persisted up to 2 μs (**Figs. 2, S1b, S2**). None of the systems reverted to active conformations within the 1–2 μs timeframe, suggesting kinetic trapping or a high reactivation barrier. Principal component analysis (PCA) of backbone atoms from 1 μs MD simulations showed that WT and F200A formed compact, unimodal clusters in PC1–PC2 space, consistent with stable active-state conformations (**Fig. 3a**). W279A and W356A sampled broader distributions with multiple sub-states, indicating increased flexibility, while double and triple mutants exhibited the greatest heterogeneity and multimodal clustering. W356A-containing mutants in particular displayed well-separated clusters, reflecting transitions between distinct conformations. Extreme-PC structures (**Fig. 3b**) revealed that WT, F200A, and W279A retained an active-like extracellular TM1, whereas W356A and all double mutants showed outward TM1 displacement characteristic of inactive states. Intracellularly, W279A, W356A, and all double mutants displayed inward TM6/TM7 movement, a hallmark of G-protein uncoupling^5,7^. The triple mutant maintained an active-like extracellular pocket but partially collapsed the intracellular cavity, consistent with an intermediate state. These results identify W356^6.4^^8^ as a central aromatic anchor in the toggle-switch network, essential for active-state stability. Its substitution disrupted intramolecular interactions, increasing conformational heterogeneity and promoting partial or full inactivation. In RMSD and PCA space, all W356A-containing mutants shifted markedly toward the inactive state, unlike F200A or W279A

**Fig. 3.**
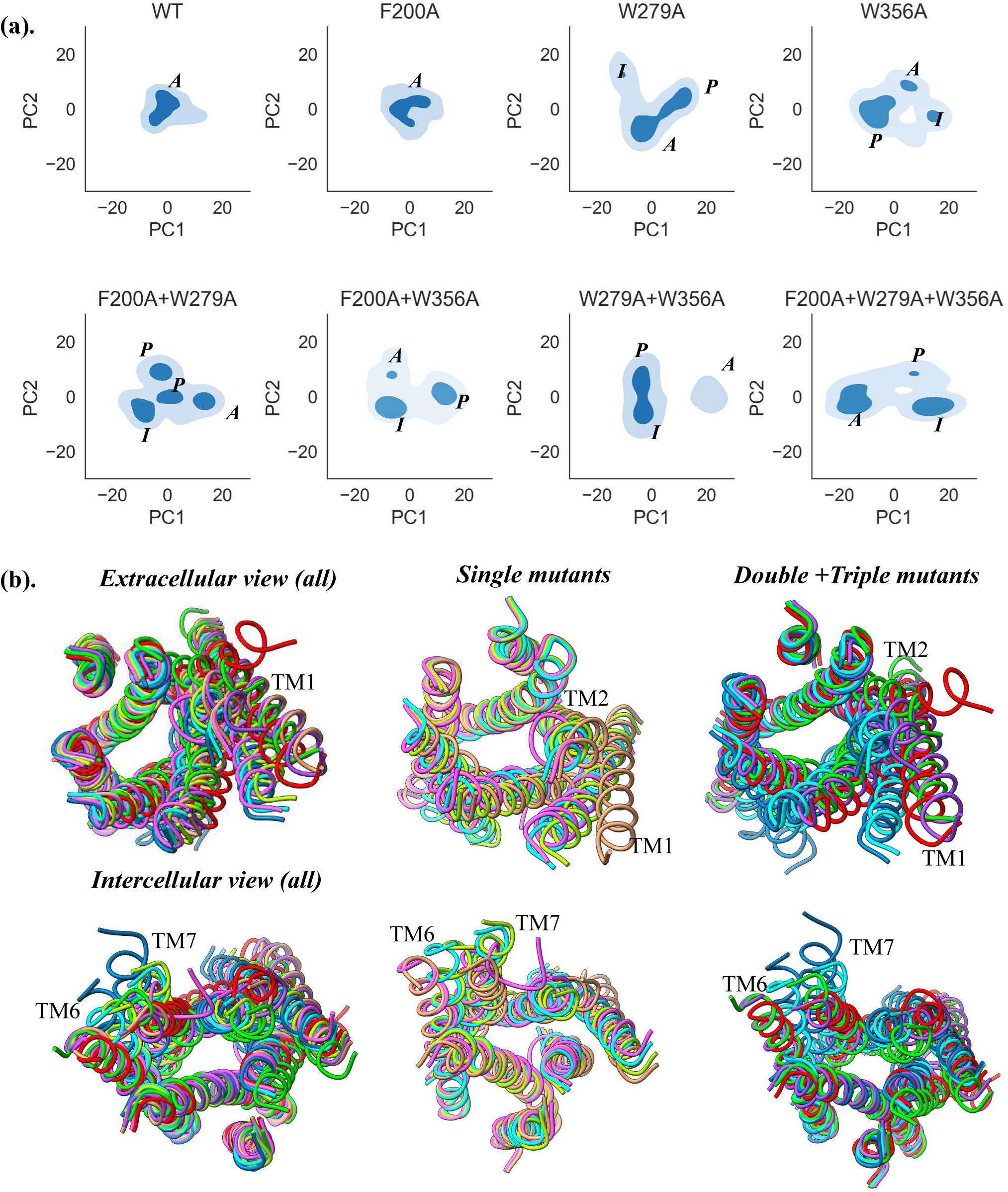
Principal component analysis reveals mutation-induced conformational heterogeneity in CB1 states. **(a)** Conformational sampling along the first two principal components (PC1 and PC2). WT shows a compact distribution of active state (*A*) population, while mutants (except F200A) exhibit broader and more dispersed ensembles including inactive states (*I*) and partial states (*P*), indicating increased flexibility. **(b)** Extreme structures along PC1 highlight structural divergence. Extracellularly, WT, F200A, and W279A retain active-like TM1 positioning, while W356A and all double mutants adopt inactive-like conformations. Intracellular views show TM6–TM7 inward collapse in W279A, W356A, and double mutants. The triple mutant preserves an active-like extracellular face but shows partial intracellular inactivation.

### Toggle switch mutations influence conserved microswitches reorganization

To explore how aromatic toggle-switch mutations influence CB1 activation, we analyzed two conserved class A GPCR motifs: the DRY (TM3) and NPxxY (TM7) switches. In canonical GPCRs, the inactive state features an ionic lock between R214^3.50^ and D338^6.30^, which is disrupted upon activation. In CB1, this lock measures 14.14 Å in the active crystal structure and 8.69 Å in the inactive state, when measured from their Cα–Cα (**Fig. 4a**)^16,27^. In our simulations, WT and single mutants (F200A, W279A, W356A) maintained large separations (>12 Å), whereas double and triple mutants, especially F200A+W356A, consistently reduced this distance (<9 Å), indicating partial ionic lock reformation and a shift toward the inactive conformation (**Fig. 4b**).

**Fig. 4:**
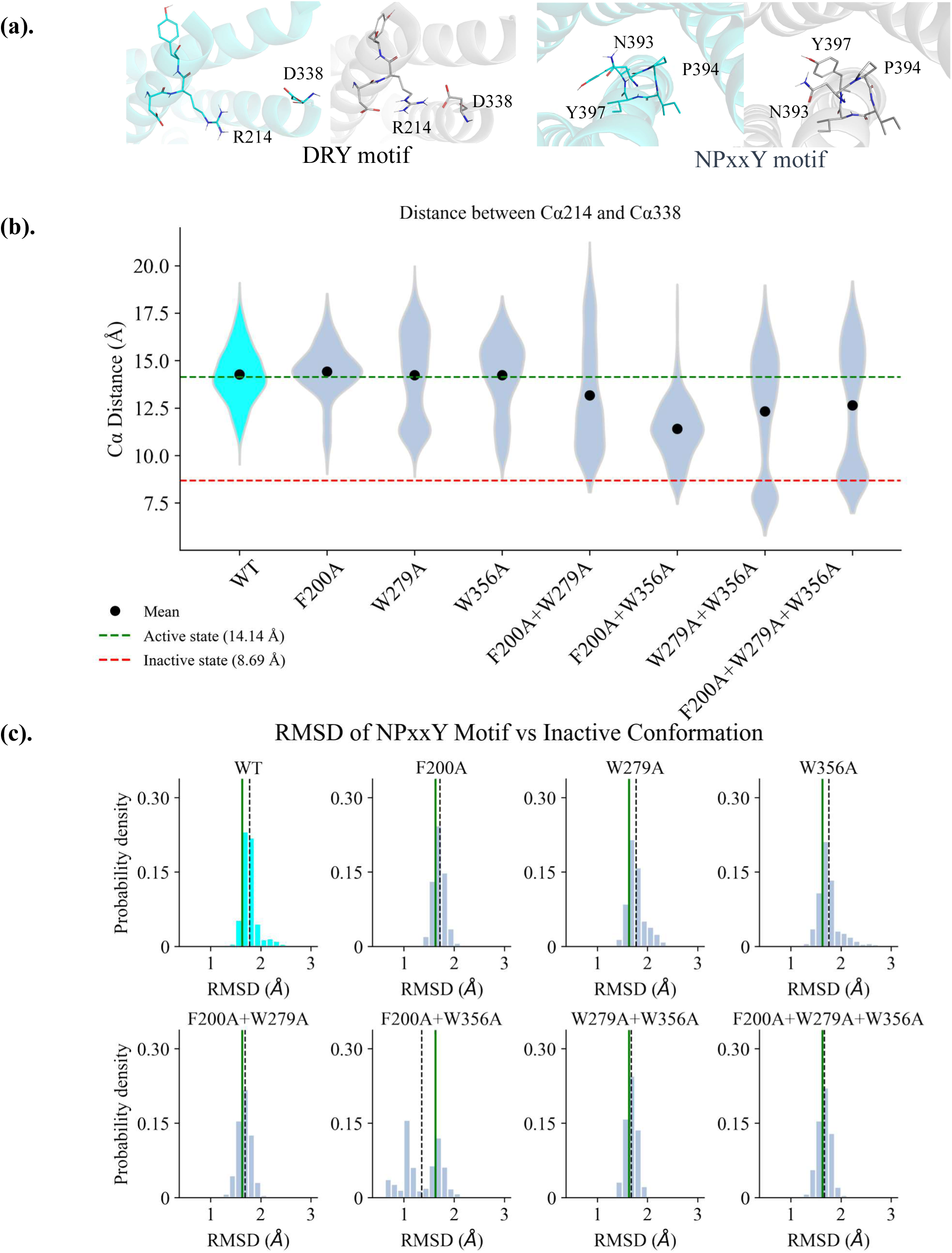
Disruption of the aromatic domain modulates conserved GPCR microswitches. **(a)** Comparison of active (cyan) and inactive (gray) state conformations of the DRY and NPxxY motifs in the CB1 receptor. These motifs, conserved across class A GPCRs, serve as molecular switches that mediate conformational transitions essential for receptor activation. **(b)** Salt bridge dynamics between R214 (DRY motif, TM3) and D338 (TM6) in inactive (red) and active (green) states, shown as Cα–Cα distances over time; black dots indicate mean values. **(c)** RMSD profiles of the NPxxY motif reveal a marked conformational shift away from the active state only in the F200A+W356A mutant. The black dotted line denotes the mean RMSD.

The NPxxY motif (N393^7.49^–Y397^7.53^) in CB1 adopts a non-canonical conformation compared to other GPCRs. In the active CB1 structure, Y397^7.53^ adopts an upward-facing rotamer, interacting with TM1 and TM2 rather than the typical TM6 engagement seen in receptors like β2AR or rhodopsin^16,17,30^. In addition, this motif plays a key role in CB1 distinctive G-protein coupling, where its mutation specifically impairs β-arrestin recruitment without affecting G-protein–mediated cAMP signaling^40^. To assess conformational changes in this motif, we calculated the RMSD of the NPxxY region relative to the inactive reference structure. Among all systems, only the F200A+W356A mutant exhibited a significant conformational drift of this motif toward the inactive state (**Fig 4c**), suggesting a potential role for the toggle switch residues in modulating NPxxY-associated activation mechanisms via altered Y397^7.53^/TM1/TM2 interactions.

Similarly, DRY motif mutations partially reduce G-protein activation, whereas double and triple mutations markedly impair G-protein signaling and enhance β-arrestin recruitment, shifting the receptor toward a β-arrestin–biased signaling profile^23^. Together, our results suggest that aromatic domains, DRY, and NPxxY motifs form a cooperative microswitch network governing CB1 conformational stability and signaling bias, with the F200A+W356A mutation uniquely positioned to destabilize the active state and reconfigure downstream coupling preferences.

### Helical reorganization in mutants supports a Seesaw model of CB1 Signaling

The activation of CB1 follows a seesaw-like motion, where the extracellular side narrows in the active state while the intracellular side opens, and the reverse occurs in the inactive state. More specifically, the seesaw movement is accomplished by the tilting of transmembrane helices TM1, TM2 and TM6^16,17,27,41^ (**Fig. 5a**). The movement of TM6 is related to G-protein coupling, while TM1 and TM2 movements are consequences of ligand binding ^16,17,27,42,43^. Thus, the state of CB1 can be characterized by the packing of TM1, TM2 and TM6. We calculated the following distances between anchor residues: TM1-TM6 on the extracellular side between Q115^1.31^ and D366^6.58^, TM3-TM6 on the intracellular side between S217^3.53^ and D338^6.30^ **(Fig. 5b)**. In the active state of CB1, the TM1–TM6 and TM3–TM6 distances measure 21.9 Å and 15.3 Å, respectively, whereas in the inactive state, these distances increase to 27.7 Å and decrease to 8.7 Å respectively. This divergence enables a clear distinction between the two conformational states within the defined structural space. Mapping the simulation trajectories onto the two-dimensional conformational space revealed that both the WT CB1 and the F200A single mutant predominantly retain active-like conformations throughout the simulations. In contrast, the W279A and W356A single mutants exhibited increased conformational flexibility, sampling a broader range of structural states. Notably, subpopulations of W279A adopted conformations with intracellular compaction, while W356A sampled conformations with extracellular widening, indicative of partial transitions toward the inactive and active states, respectively. These effects were further amplified in the double mutants, particularly in the F200A+W356A combination, which displayed a pronounced shift toward the inactive state. The triple mutant (F200A+W279A+W356A) sampled a substantial ensemble of conformations corresponding to a ‘partially active’ state, characterized by simultaneous compaction at both the extracellular and intracellular regions, reflecting a distinct intermediate along the activation–inactivation continuum.

**Fig. 5.**
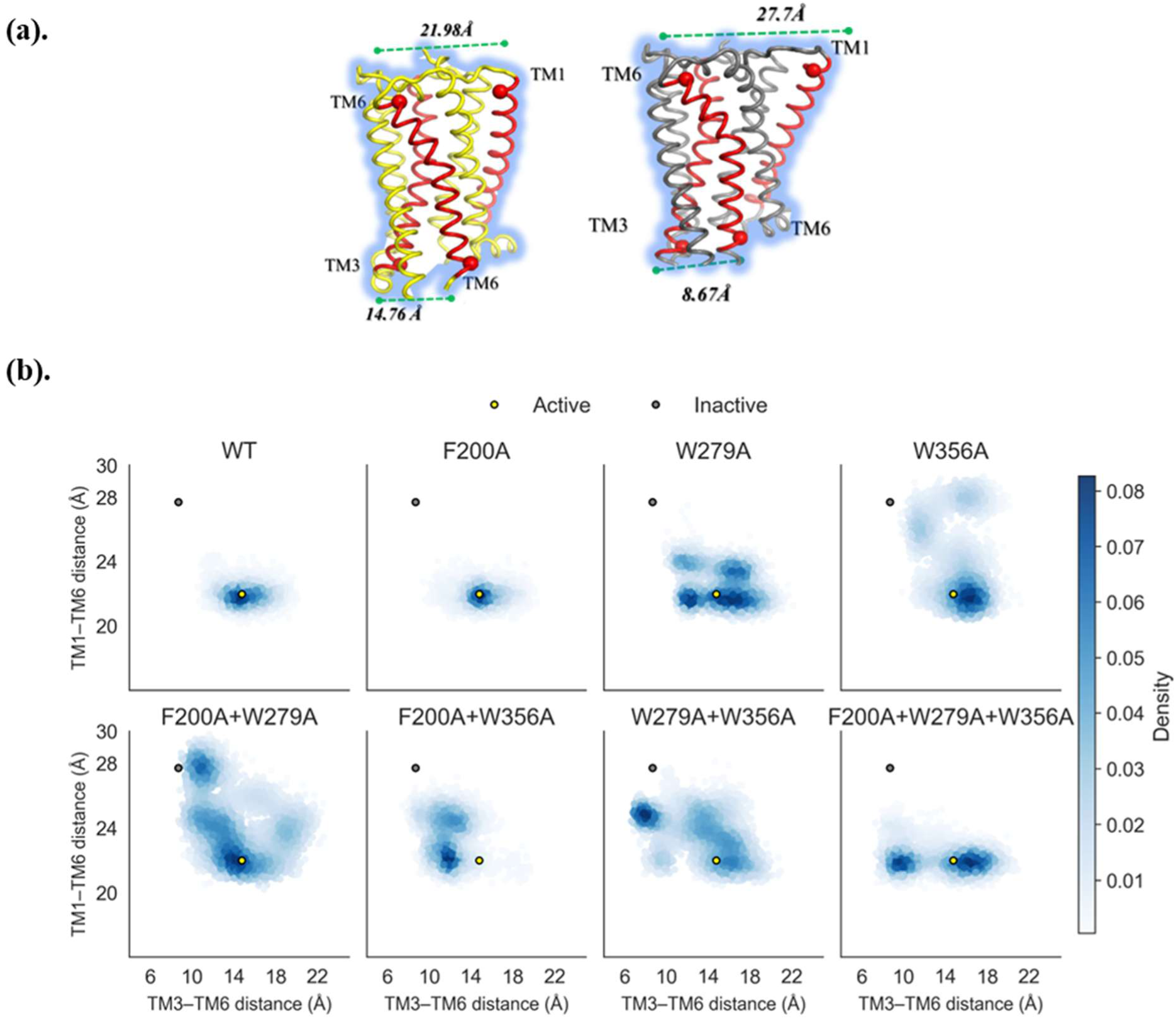
Conformational dynamics of CB1 transmembrane region. **(a)** Active (yellow) and inactive (gray) conformation structures of CB1, with Cα atoms of residues Q115 of TM1, D366 of TM6 (extracellular) and S217 of TM3 and D338 of TM6 (intracellular) annotated to highlight key helical movements in activation. **(b)** Distances of TM1–TM6 (extracellular) vs. TM3–TM6 (intracellular) movements from simulations. Reference distances from active and inactive states are shown in yellow and gray points, respectively.

To better illustrate helix-specific changes, we computed 7×7 RMSD matrices comparing each TM helix across all systems to the inactive-state structure. Unlike global TM bundle RMSD, each 7×7 RMSD matrix was generated by iteratively superposing the structure on one TM helix (e.g., TM1) and computing the backbone RMSD for TM1–TM7, producing a matrix of localized RMSD values. This method accounts for internal rearrangements while minimizing the confounding effects of rigid-body rotations or global shifts^44^. The matrices were calculated for the most populated conformations of each system (i.e., the largest cluster structures) and the minimum-RMSD structures (i.e., the most inactive-like conformers, based on the lowest global TM backbone RMSD to the inactive-state structure). The RMSD results confirm that alanine substitutions at F200, W279, and W356 destabilize the active-state conformation in the absence of a ligand, shifting helix orientations toward inactive-like states. Notably, the triple mutant (F200A+W279A+W356A), double mutant F200A+W356A, and single mutant W356A exhibited cluster and minimum structures with helix orientations closely resembling the inactive state, as evidenced by small RMSD values in their matrices (**Fig. S3**). The consistent presence of W356A in these mutants suggests that W356 may play a critical role in stabilizing the active-state helical arrangement, potentially acting as a key allosteric gatekeeper. However, the comparable inactive-like conformations in the triple and F200A+W356A mutants indicate cooperative effects among F200, W279, and W356, warranting further quantitative analysis to confirm the dominant influence of W356.

### The roles of residue F200^3.36^, W279^5.43^ and W356^6.48^

Structural models from the minimum RMSD frames (**Fig. S5**) showed that, on the extracellular side, TM1 and TM2 remained in an active-like packed conformation in the WT and F200A systems. In contrast, the TM1 in W356A and F200A+W356A mutants clearly shifted away from the packed conformation, consistent with the overall movement toward an inactive-like state. On the intracellular side, TM6 underwent a shift toward its inactive orientation in all W356A-containing double and triple mutants. In contrast to F200^3.36^ and W356^6.48^, which directly controlled intracellular microswitches and active-state stability, W279^5.43^ contributed primarily to extracellular pocket organization. In our simulations, W279A promoted partial inactivation while maintaining overall cavity architecture, consistent with experimental data showing that this mutation markedly reduces binding of the antagonist Rimonabant by disrupting aromatic stacking interactions with the TM3–TM5–TM6 aromatic microdomain^29^. This suggests that W279^5.43^ functions less as a toggle switch and more as a structural anchor, stabilizing ligand orientation and extracellular pocket integrity during conformational transitions.

While representative structures from the most populated clusters reflect more moderate deviations (**Fig. S6**), they reinforce the trend toward inactivation. These differences should be interpreted in light of two factors: (1) the “most inactivated” structures reflect extreme conformations sampled during unbiased MD simulations, whereas the representative models capture average ensemble behavior; and (2) the simulation timescale (∼1 μs) may be insufficient to capture complete inactivation transitions. Taken together, our structural data underscore the progressive and mutation-specific destabilization of the active state and highlight the key role of the aromatic domain in maintaining conformational integrity in the absence of ligand. In this context, W356^6.48^ serves as a structural pivot, F200^3.36^ as a kinetic gate, and W279^5.43^ as a modulatory buffer that stabilizes the orthosteric geometry and tunes the receptor conformational response to perturbation.

### Targeted MD simulations reveal energetic and structural barriers to CB1 inactivation

To further probe the energetic barriers associated with CB1 inactivation, we used targeted MD simulations starting from the active-state conformation towards the inactive state. When the restraining force was applied solely to the Cα atoms of TM helices, TM rearrangements progressed toward the inactive state at gradually increasing force constants (0.1, 0.2, 0.4, and 0.8 kcal·mol⁻¹·Å⁻²), yielding RMSD values of 1.994 Å, 1.318 Å, 0.888 Å, and 0.617 Å, respectively, compared to the inactive state structure. However, in all cases, F200 retained its active-state rotamer, suggesting that TM repacking and toggle switch transition are decoupled, with the latter requiring specific triggers, such as ligand binding (**Fig. S7a**).

When the force was applied to all heavy atoms in TM helices, the energetic requirement for inactivation decreased. At a low force constant (0.01 kcal·mol⁻¹·Å⁻²), TM rearrangement was initiated in several trajectories, with RMSD values approaching ∼1.75 Å when compared to the inactive conformation; however, F200 still remained in an active-like conformation. Complete inactivation, including F200^3.36^ rotamer flipping, was observed only at higher force constants (0.05 kcal·mol⁻¹·Å⁻²), with F200 transitioning to its inactive orientation once the TM RMSD approached ∼0.76 Å (**Fig. 7a-b**). These findings demonstrate that the toggle switch represents a distinct energetic hurdle, beyond global TM reorganization.

**Fig. 6.**
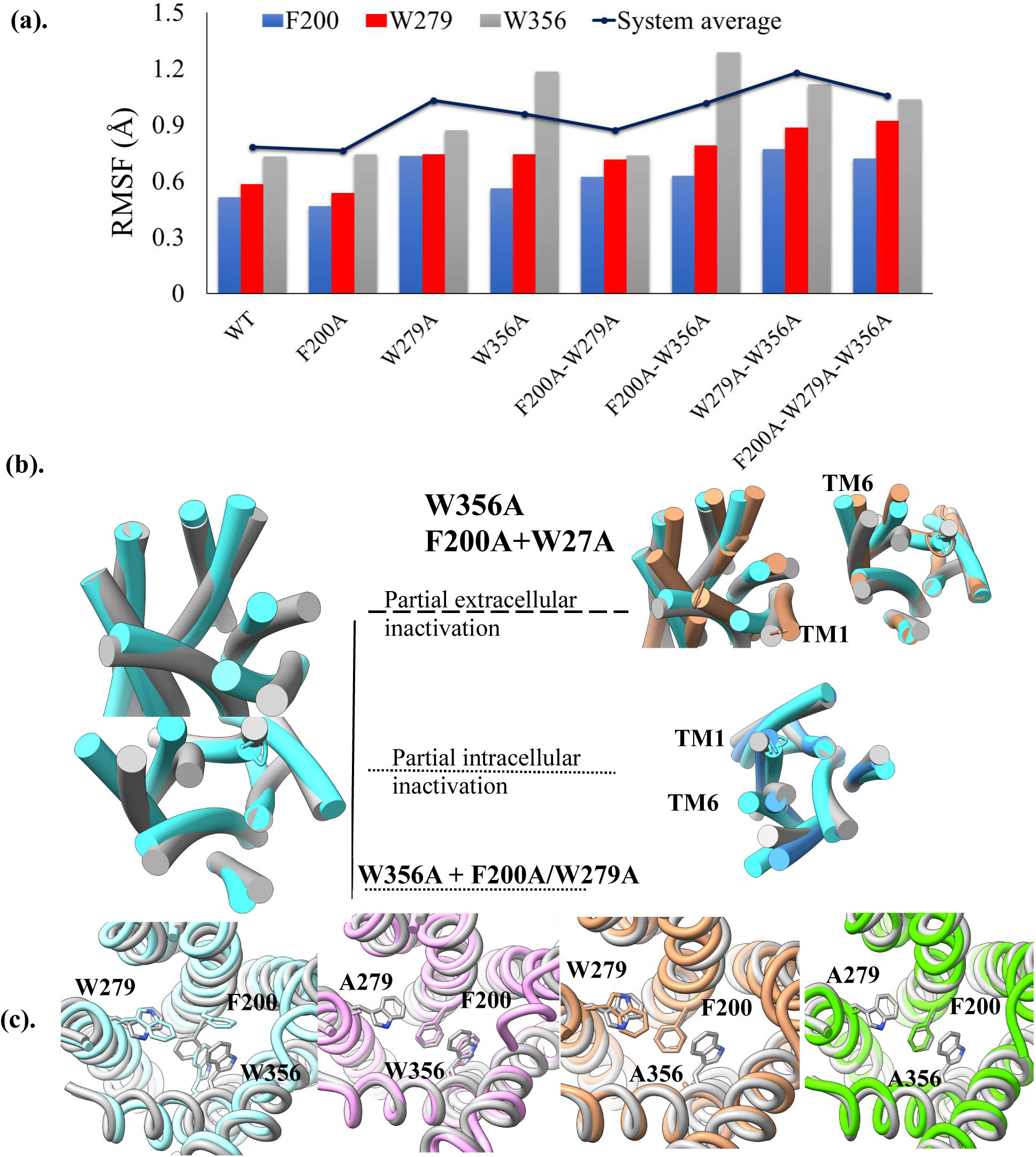
Mutations in the aromatic toggle domain alter local flexibility and rotameric switching. **(a)** Backbone RMS fluctuations of the mutated residues (F200, W279, W356) for each system, computed from the last 500 ns of simulation (500–1000 ns). The system-wide average RMSF is shown as a reference line. Notably, mutations at W356^6.48^ consistently lead to elevated fluctuations at this position, suggesting increased conformational plasticity. **(b)** Superimposition of the snapshot from simulations exhibiting partial activation (extracellular or intracellular) onto the reference crystal structures of the active (cyan) and inactive (gray) CB1 receptor. The W356A mutant dominantly demonstrates partial extracellular inactivation, whereas intracellular partial inactivation is more prominent in mutants combining W356A with F200A and/or W279A. **(c)** Representative toggle switch conformations from frames most similar to the inactive state (i.e., lowest RMSD), highlighting conformational shifts of F200^3.36^ in response to alanine mutations at W279^5.43^ and W356^6.48^.

**Fig. 7.**
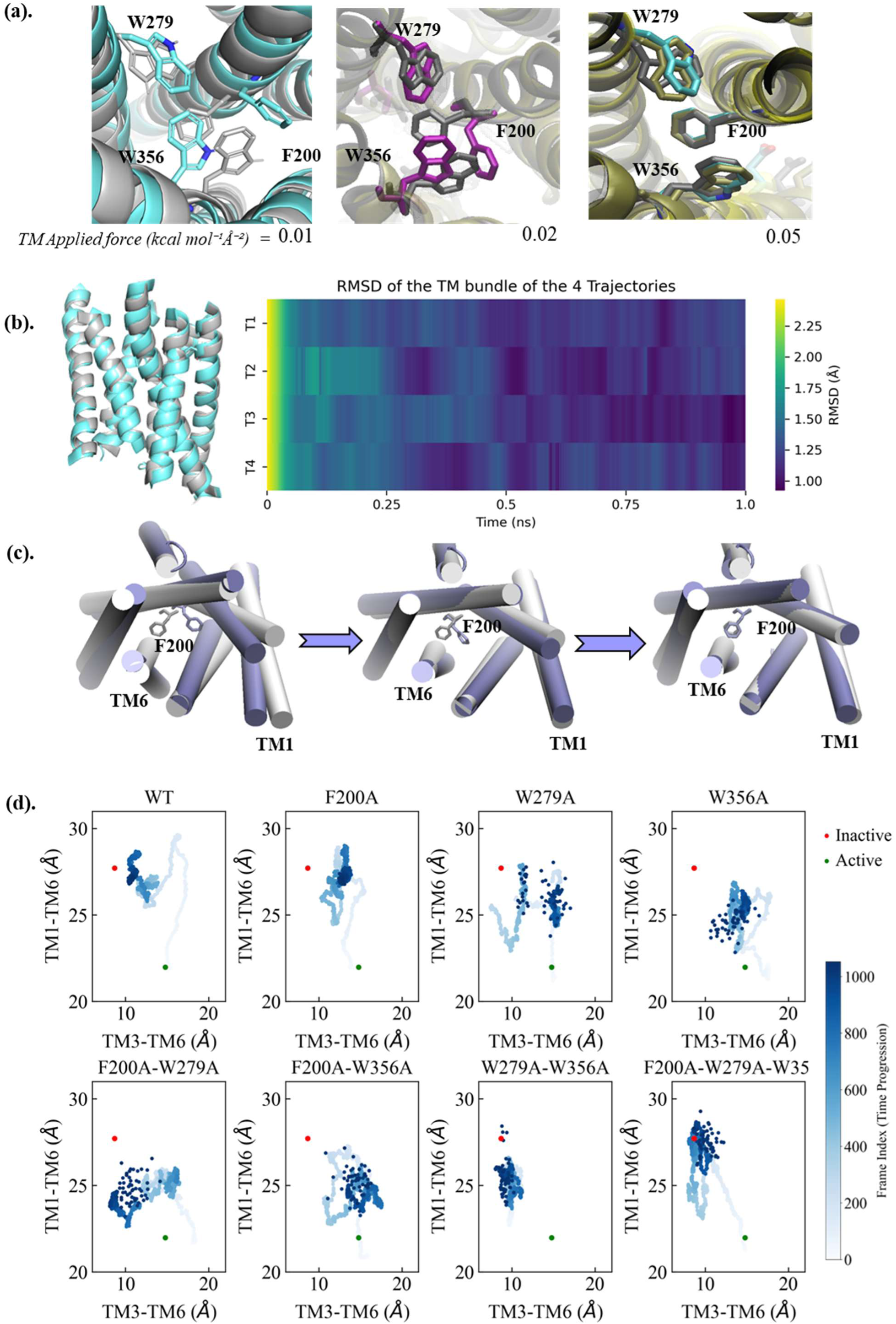
Targeted MD simulations reveal force-dependent inactivation and sequential conformational transitions in CB1. **(a)** Structural superposition of the experimental inactive CB1 structure (gray) with final frames from tMD simulations applying forces of 0.01, 0.02, and 0.05 kcal·mol⁻¹·Å⁻². Only the highest force (0.05) drives F200^3.36^ into its inactive-state rotamer, highlighting the elevated energetic barrier associated with toggle switch flipping. **(b)**. Time evolution of TM bundle RMSD in WT CB1 during steered MD simulations applying a force of 0.01 kcal·mol⁻¹·Å⁻² to all TM atoms. **(c)** Extracellular compaction precedes intracellular gate closure and F200 reorientation, illustrating a sequential inactivation mechanism in which toggle switch flipping is a high-barrier event. **(d)** TM1–TM6 (Q115^1.31^– D366^6.58^) and TM3–TM6 (S217^3.53^–D338^6.30^) Cα–Cα distances measured during tMD simulations under 0.01 kcal·mol⁻¹·Å⁻² force. Both WT and F200A systems exhibit early extracellular compaction followed by delayed intracellular rearrangement, consistent with stepwise progression toward the inactive state.

Tracking helical rearrangements using distances between Q115^1.31^ and D366^6.58^ (extracellular TM1–TM6) and between S217^3.53^ and D338^6.30^ (intracellular TM3–TM6) revealed that extracellular widening precedes intracellular compaction and F200 reorientation (**Fig. 7c**). These results support a sequential mechanism for inactivation, where toggle switch flipping is a late-stage, high-barrier event.

### Disruption of the aromatic domain lowers the energetic barrier and alters the CB1 inactivation pathway

To evaluate the role of the F200^3.36^, W279^5.43^, and W356^6.48^ residues in stabilizing the active conformation of CB1, targeted MD simulations were also performed for mutant systems with alanine substitutions at these positions. A constant force of 0.01 kcal·mol⁻¹·Å⁻² was applied to all transmembrane (TM) atoms. Remarkably, this minimal perturbation was sufficient to drive all mutant systems into inactive-like conformations (**Fig. 7d)**, with significantly lower RMSD values compared to the wild-type receptor under the same conditions (**Fig. S7b**). Notably, in all systems where F200 remained in its wild type form (i.e., W279A, W356A, and W279A + W356A), we observed spontaneous rotameric flipping of the F200 side chain into its inactive conformation. This supports the hypothesis that the π–π interactions within the aromatic domain contribute to the energetic barrier maintaining the active state (**Table 2**). Similar to the WT, F200A mutant inactivation pathway proceeds sequentially initial extracellular widening (TM1–TM6 distance increase) followed by intracellular TM6 closure and F200 rotamer flipping (**Fig. 7c**).

**Table 1:**
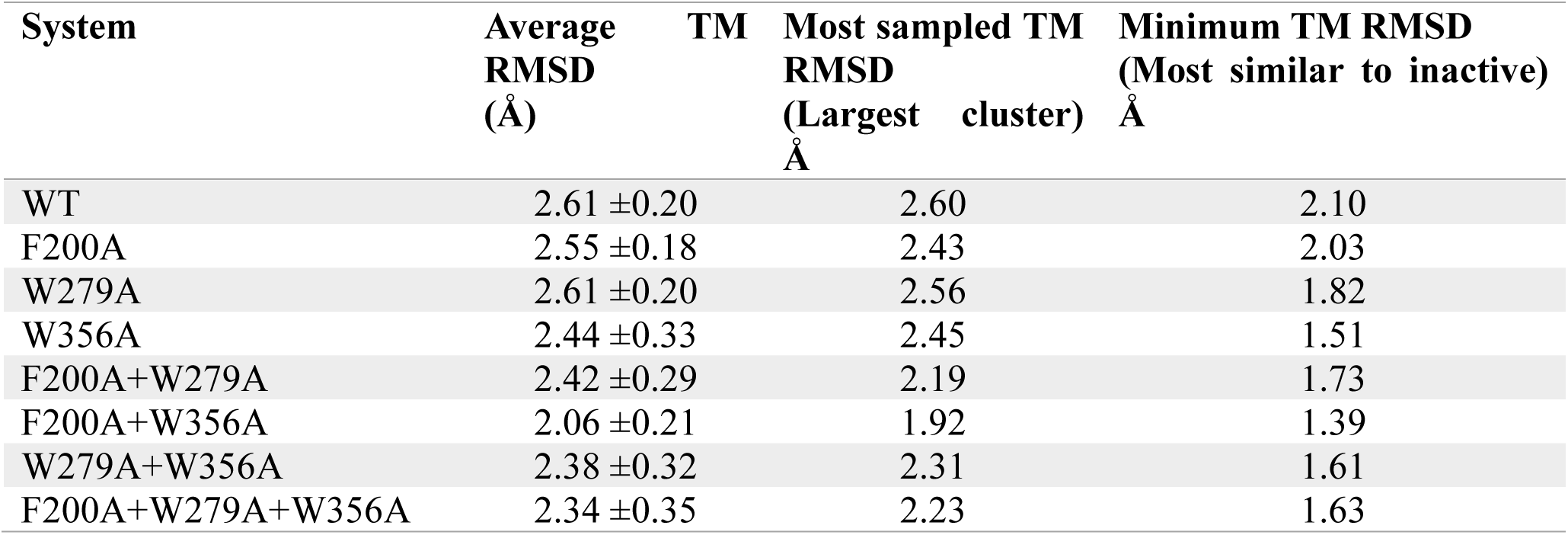
Representative structures from the equilibrium MD simulation.

**Table 2:** Representative structures from targeted MD simulation (TM applied force= 0.01Kcal/M/A)

| System | Average RMSD<br>(TM BB)<br>Å | Minimum RMSD<br>(TM BB)<br>Å | RMSD (TM All<br>atoms)<br>Å |
| --- | --- | --- | --- |
| WT | 1.272 ±0.238 | 1.008 | 1.100 |
| F200A | 1.218 ±0.273 | **0.937** | 1.034 |
| W279A | 1.453 ±0.252 | **1.005** | 1.423 |
| W356A | 1.338 ±0.303 | **0.993** | 1.209 |
| F200A+W279A | 1.386 ±0.281 | **0.944** | 1.338 |
| F200A+W356A | 1.416 ±0.251 | **1.014** | 1.415 |
| W279A+W356A | 1.284 ±0.296 | **0.912** | 1.228 |
| F200A+W279A+W356A | 1.281 ±0.331 | **0.866** | 0.986 |

However, the inactivation pathways diverged significantly between variants. While the WT and F200A mutants followed the canonical sequential pathway with extracellular (TM1-TM2) widening preceding intracellular (TM6) rearrangement, single W279A and W356A mutants showed an attenuated pathway whereby the rearrangements occur earlier, with a reduced extent of extracellular widening observed. Most notably, double (W279A+W356A) and triple mutants achieved the most inactive-like conformations (minimum RMSD 0.912 Å and 0.866 Å, respectively) through concerted extracellular and intracellular transitions, suggesting complete disruption of the “gate”. This pathway alteration likely stems from both steric (removal of bulky tryptophan side chains) and energetic (reduced torsional barriers) effects, as evidenced by mutants reaching inactive states faster than the wild type under identical conditions (**Fig. 7d**). These results collectively demonstrate that the aromatic domain serves as a dual steric and energetic gate controlling both the magnitude and temporal coordination of the CB1 inactivation pathway.

## Discussion

MD simulations highlight the critical role of the aromatic toggle switch domain (F200^3.36^, W279^5.43^, W356^6.48^) in stabilizing the active conformation of CB1 and controlling its transition to the inactive state. In WT CB1, strong π–π interactions stabilize the core TM bundle and restrict conformational fluctuations. Alanine mutations disrupt aromatic interactions, increasing local flexibility and promoting inactivation. In particular, the W356A and F200A+W356A mutants exhibited both structural instability and pronounced movements of TM1 and TM6, mimicking features of the inactive state. Interestingly, the single F200A mutant retained a WT-like conformation over 1 μs of simulation, but in systems where F200^3.36^ remained native and W356^6.48^ was mutated, the F200^3.36^ sidechain readily adopted an inactive-like rotamer. This implies that the π–π stacking interaction between F200^3.36^ and W356^6.48^ is essential not only for maintaining the active state but also for preventing premature inactivation. These findings support a functional division within the aromatic domain: W356^6.48^ acts as a structural pivot whose integrity maintains orthosteric packing, F200^3.36^ serves as a kinetic gate that resists conformational switching, and W279^5.43^ functions as a modulatory buffer, preserving orthosteric architecture and tuning the conformational response to perturbations.

These structural transitions align with previous biochemical studies. For instance, F200A has been shown to enhance the constitutive activity of CB1 receptor^20^, likely by disrupting the stabilizing lock (π-stacking with W356^6.48^) that prevents spontaneous activation. Likewise, W279A compromises antagonist binding by disrupting aromatic stacking interactions^29^, aligning with our observation that this mutation weakens extracellular pocket stability and promotes partial inactivation without collapsing the cavity.

Targeted MD simulations further reveal that rotamer switching of F200^3.36^ is a high-barrier process that requires greater energy input than TM helical rearrangement. This distinction was evident in WT systems, where TM movements occurred readily but F200^3.36^ remained in the active-state rotamer. In contrast, mutations in the toggle switch domain lowered the energy barrier, allowing inactivation at minimal force inputs. These findings provide a mechanistic framework for understanding the structural determinants of CB1 conformational transitions. The toggle switch domain functions as a molecular clutch that controls activation state transitions; disruption of the structural pivot (W356^6.48^) promotes collapse of the active conformation, while mutation of the kinetic gate (F200^3.36^) further lowers the barrier to inactivation in a cooperative manner. From a drug discovery perspective, this offers new avenues for selectively modulating CB1 activity by targeting these aromatic interactions. Inverse agonists or allosteric modulators designed to destabilize this network may enhance inactivation, while stabilizers of the aromatic core could preserve active-state signaling.

While our simulations offer critical insights, they also highlight the limitations of accessible timescales. The observed transitions toward inactivation occurred within 1 μs, but full equilibrium between active and inactive states was not reached. Nevertheless, the directional trends such as TM6 inward motion and F200^3.36^ rotamer flipping strongly suggest an inactivation pathway initiated by domain destabilization. Future enhanced sampling simulations or experimental validation could further clarify the energy landscape and intermediate states involved.

### Physiological implications of CB1 inactivation bias

Our simulations demonstrate that alanine substitutions within the aromatic domain bias the receptor toward inactive-like conformations, even in the absence of ligands, reducing its constitutive activity. This conformational shift has significant physiological implications for central nervous system (CNS) and peripheral functions, underscoring the gating role of the aromatic domain. In CNS, CB1 modulates synaptic transmission through retrograde inhibition of GABA and glutamate release^45^. Constitutive activity sustains basal endocannabinoid tone, stabilizing synaptic balance and reducing neuronal excitability^46^. Mutations favoring inactivation such as F200A+W356A may disrupt this tone, increasing excitability and potentially impairing memory consolidation, stress adaptation, and emotional regulation^47–49^. Additionally, CB1 receptor modulation of mesolimbic dopaminergic signaling suggests that reduced activity could contribute to anhedonia^50,51^ or depressive-like behaviors, as observed with the inverse agonist rimonabant^52^.

Peripherally, CB1 regulates metabolic and inflammatory processes in the liver, adipose tissue, and immune cells. Attenuated CB1 signaling, as induced by mutations, reduces appetite, enhances insulin sensitivity, and promotes anti-inflammatory response ^53–56^. These effects support therapeutic potential of CB1 in obesity, liver fibrosis, and metabolic syndrome. However, chronic inactivation may compromise protective endocannabinoid signaling under physiological stress, potentially exacerbating conditions like epilepsy or chronic pain^57^. The tunable inactivation bias induced by aromatic domain mutations provides mechanistic insights into the CB1 activation landscape. Targeting the aromatic toggle switch could enable the design of selective therapeutics that dampen excessive basal signaling while preserving stimulus-dependent activation, mitigating adverse effects associated with complete CB1 antagonism^17^. Such modulators could reduce basal CNS endocannabinoid tone and enhance metabolic control, beneficial for obesity, but may pose risks in neurological or psychiatric disorders where endocannabinoid signaling supports neuroprotection and emotional regulation^45,57^.

In summary, mutations biasing CB1 toward inactivity mimic pharmacological antagonism, with implications for metabolic regulation, CNS excitability, and endocannabinoid signaling (**Fig. S8**). These findings offer a structural and functional framework for developing CB1 modulators with improved therapeutic precision.

## Conclusion

Using all-atom MD simulations, we demonstrate that the aromatic domain comprising F200^3.36^, W279^5.43^, and W356^6.48^ functions as a conformational lock that stabilizes the active state of the CB1 receptor. These residues form a tightly packed aromatic domain within the orthosteric site and are central to the toggle-switch mechanism that regulates CB1 constitutive activity. Single-point alanine substitutions at these positions introduce only minor local perturbations; however, double and triple mutations, particularly F200A+W356A, substantially disrupt the structural integrity of the receptor, promoting transitions toward inactive-like conformations.

Equilibrium MD simulations revealed a progressive increase in conformational flexibility and helical rearrangement with increasing mutation load, consistent with a cumulative “trimming effect” that weakens π–π interactions and loosens TM helix packing. The residue RMS fluctuations, principal component analysis, and transmembrane helix analyses identified W356^6.48^ as a key structural gatekeeper maintaining active-state topology, while targeted MD simulations revealed that F200^3.36^ acts as a high-energy rotameric barrier, only flipping to its inactive state under elevated force input. Notably, toggle switch mutations also modulated the conformations of the DRY and NPxxY motifs, two conserved GPCR microswitches, implying a cooperative network that tunes signaling bias.

Targeted MD of mutant systems revealed that disruption of the aromatic domain alters the sequence and energetics of the inactivation pathway, enabling inactivation at minimal force levels. These findings provide mechanistic insight into how the aromatic domain modulates the conformational ensemble of CB1, functioning as a structural rheostat that tunes signaling tone. The ability to bias CB1 toward inactive or partially active states by modulating this domain has direct implications for therapeutic design. Rather than fully antagonizing CB1, an approach associated with adverse neuropsychiatric effects, future strategies might exploit the toggle switch to selectively modulate basal activity. Such approaches could yield safer, more targeted treatment options for conditions such as pain, metabolic syndrome, and psychiatric disorders.

## Acknowledgements

This work was supported by the National Key Research and Development Program of China (2021YFA0911100), the Strategic Priority Research Program of the Chinese Academy of Sciences (XDB0480000), the National Natural Science Foundation of China (32170672 and 32000886), the Guangdong Basic and Applied Basic Research Foundation (2021A1515012461), the Shenzhen Science and Technology Program (ZDSYS20220606100606013), and the Guangdong Grants (2021QN02Y554). We thank Prof. Suwen Zhao of the iHuman Institute of ShanghaiTech University for her helpful discussions and guidance.

## Methods

### Equilibrium MD Simulation

The active (PDB: 5XRA)^16^ and inactive conformation (5TGZ)^27^ of CB1 were obtained from the Protein Data Bank (PDB)^58^. Flavodoxin inserted in ICL3 was removed, and the loop was reconstructed using Modeller v9.18^59^. In the active conformation, alanine mutations targeting residues F200^3.36^, W279^5.43^ and W356^6.48^ were introduced in all possible combinations to create CB1 mutants. As a result, seven mutant receptors containing single, and double (F200A+W279A, F200A+W356A, and W279A+W356A) and triple (F200A+W279A+W356) mutations were generated. The protonation and tautomeric states of Asp, Glu, Arg, Lys, and His residues were adjusted according to their correct charges at pH of 7.0.

The Positioning of Proteins in Membranes (PPM)^60^ server was used to orient the systems, ensuring that the receptor is properly embedded in the POPC (1-palmitoyl-2-oleoyl-sn-glycero- 3-phosphocholine) lipid bilayer. The oriented proteins were inserted in POPC bilayer using the CHARMM-GUI Membrane Builder tool^61^. In addition to the protein complex, POPC molecules, TIP3P water molecules, and excess sodium chloride ions were added to maintain an ion concentration of 0.15M, each system contained approximately 65,000 atoms. The minimum distance from protein atoms to boundaries of water box was 12 Å. The systems were modelled using CHARMM36 forcefield parameters^62^.

Before system equilibration, the steepest descent algorithm was used to minimize each system. The first stage of equilibration was for 1 ns in the NVT ensemble conditions (constant particle number, volume, and temperature). The second stage of equilibration consisted of an additional 30 ns NPT ensemble simulation (constant particle number, pressure, and temperature) with a gradual decrease in force constant restraints (from 10 to 0 kcal/mol· Å²). The force restraints applied included harmonic restraints to the heavy atoms of the protein and planar restraints to hold the position of lipid head groups of membranes along the Z-axis. Once all the equilibration steps were completed, the constraints were removed, and all systems were propagated under NPT conditions for 1000 ns with a time step of 2 fs at 310 K maintained using a Nosé-Hoover thermostat^63,64^, with a coupling time constant of 1.0 ps and pressure of 1 atm using semi-isotropic coupling^65^. MD simulations were carried out using GROMACS 5.1.2^66^. For each system, three replicates were generated by randomizing initial velocities, totaling simulation time to 24 μs.

Trajectory analyses such as RMSD (Root-mean-square deviation), RMSF (Root-mean-square fluctuation), and clustering were performed by the analysis tools in the GROMACS 5.1.2 package and VMD^67^. For structural and dynamic analyses, snapshots of each system were taken every 1 ns. Additional post-processing analyses of the trajectories, such as principal component analysis (PCA) were carried out using the MDAnalysis library package^68^. TM bundle RMSD was computed after aligning their backbone atoms to reference active and inactive structures. The TM bundle helices were defined as residues 114 to 143 for TM1, 154 to 178 for TM2, 186 to 219 for TM3, 230 to 253 for TM4, 273 to 304 for TM5, 339 to 367 for TM6 and 373 to 400 for TM7.

### Targeted MD Simulation

Targeted molecular dynamics (tMD) simulations were conducted using the AMBER18 software^69^ with the ff14SB force field^70^ to study conformational transitions toward the inactive CB1 conformation. The active CB1 receptor (∼5,100 atoms) was simulated using the generalized Born implicit solvent model with modified Bondi radii, and a 12 Å cutoff for non-bonded interactions during minimization and equilibration steps^71^. In the production phase, an infinite non-bonded cutoff (999.99 Å) was used with no periodic boundary conditions, and the system was maintained at 300 K using a Berendsen thermostat^72^. tMD was applied to guide the CB1 receptor toward an inactive conformation by imposing harmonic restraints on specific regions, including alpha carbons (Cα) of transmembrane (TM) residues or all TM atoms. As with equilibrium MD simulations, 8 systems were simulated including WT and variants with alanine mutations in the aromatic domain (F200^3.36^, W279^5.43^ and W356^6.48^). Force constants ranging from 0.01 to 0.8 kcal·mol⁻¹·Å⁻² were tested across multiple trajectories to ensure reproducibility. Conformational changes in TM helices and F200 rotamer states were analyzed using AmberTools18^73^.

