## supplemental figures for "Aromatic amino acids in the orthosteric region regulate cannabinoid receptor 1 conformation transitions"

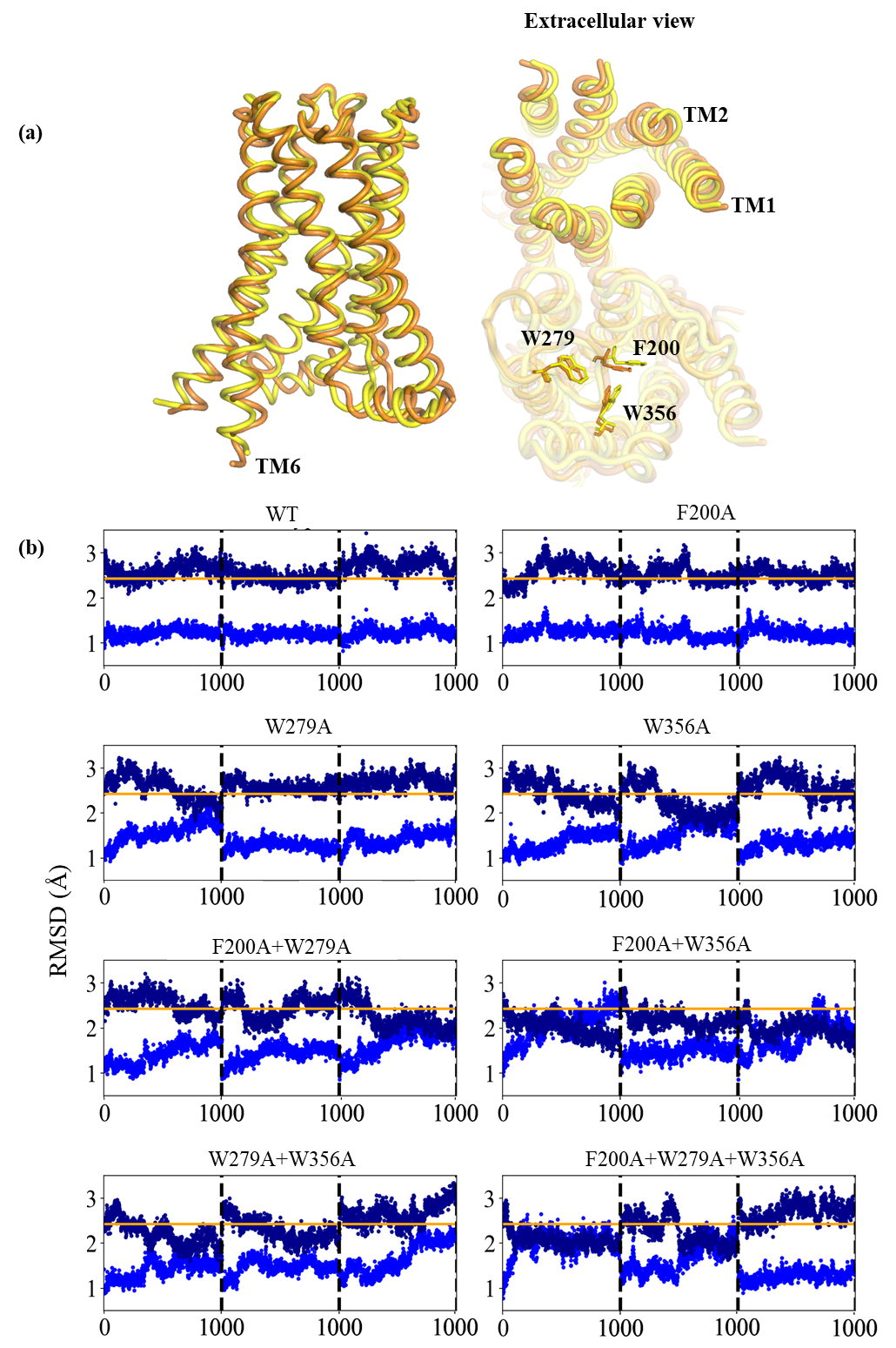
**Figure S1**: **Global conformation differences**. **(a)** MD simulation of WT CB1 initiated from the active crystal structure (yellow) shows minimal deviation, with transmembrane (TM) backbone RMSD not exceeding 0.66 Å (orange). The toggle switch residues F200 and W356 remained in their active conformations throughout the simulation. **(b)** Time evolution of TM region RMSD relative to the active (blue) and inactive (dark blue) crystal structures over 1000 ns. The orange line denotes the structural RMSD between the active and inactive TM backbones, providing a reference for the conformational landscape sampled.

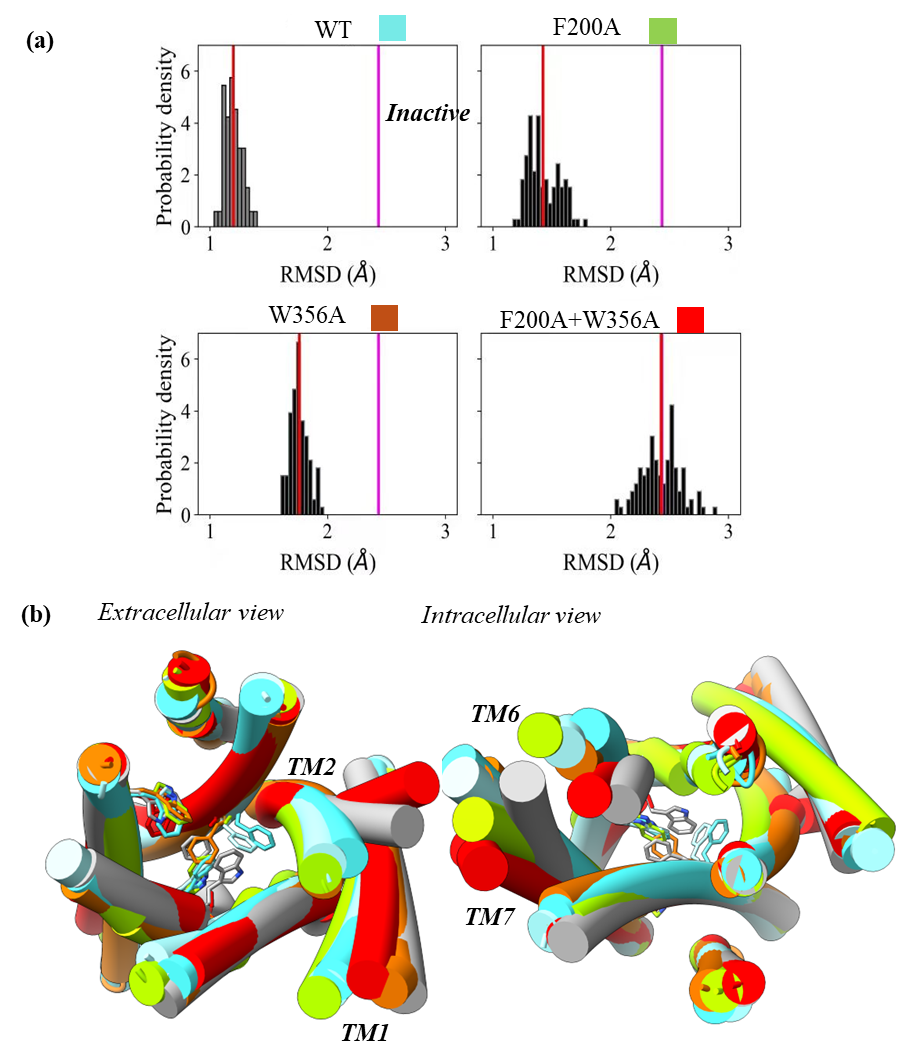

**Figure S2. Toggle switch mutations drive CB1 inactivation**. **(a)** RMSD profiles from the extended MD simulations (2000 ns) for toggle switch mutants relative to the inactive CB1 structure reveal distinct effects on conformational stability. The F200A mutation alone had minimal impact, W356A led to partial inactivation, while the combined F200A+W356A mutation resulted in complete inactivation, a disruption of the active state. **(b)** Final structures from each trajectory superposed onto the experimental inactive (gray) and active (cyan) conformations, illustrating the extent of conformational convergence.

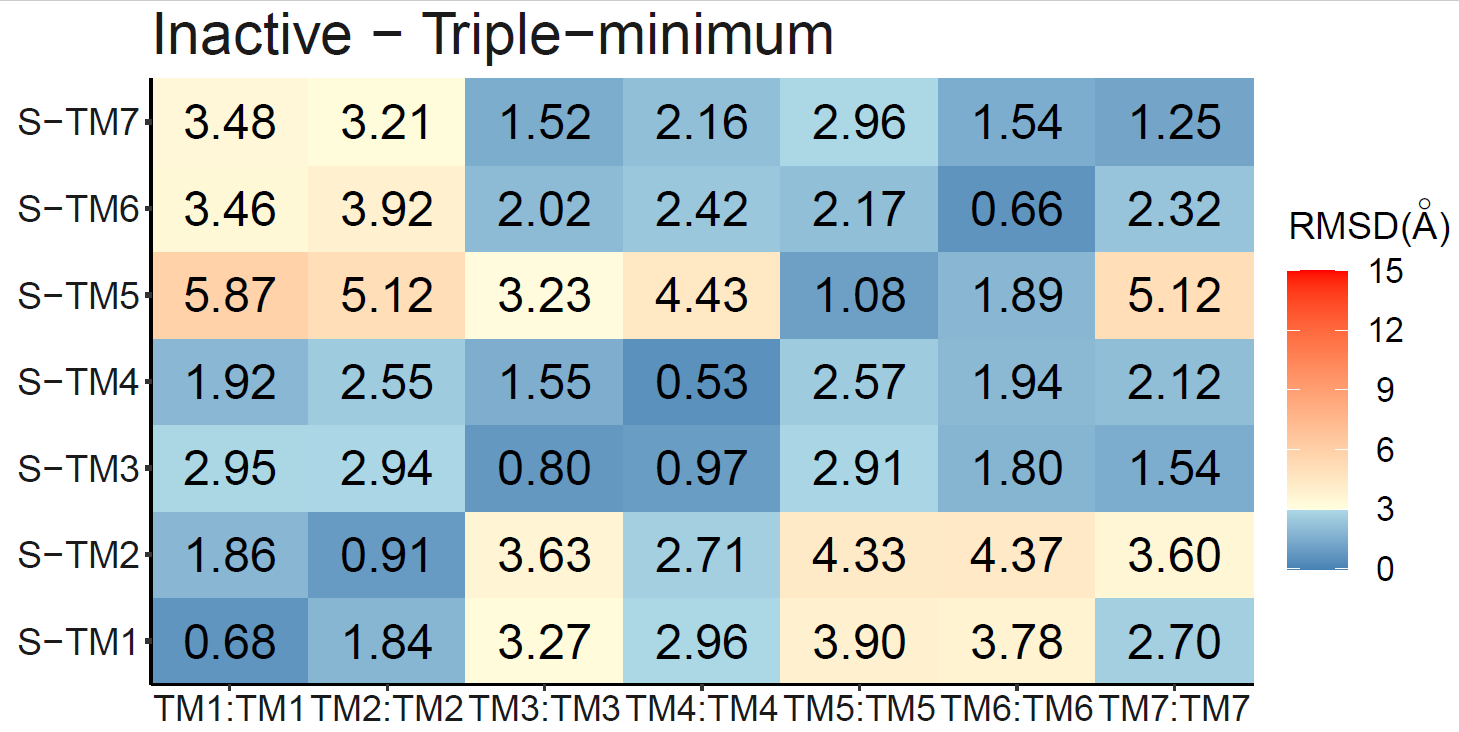

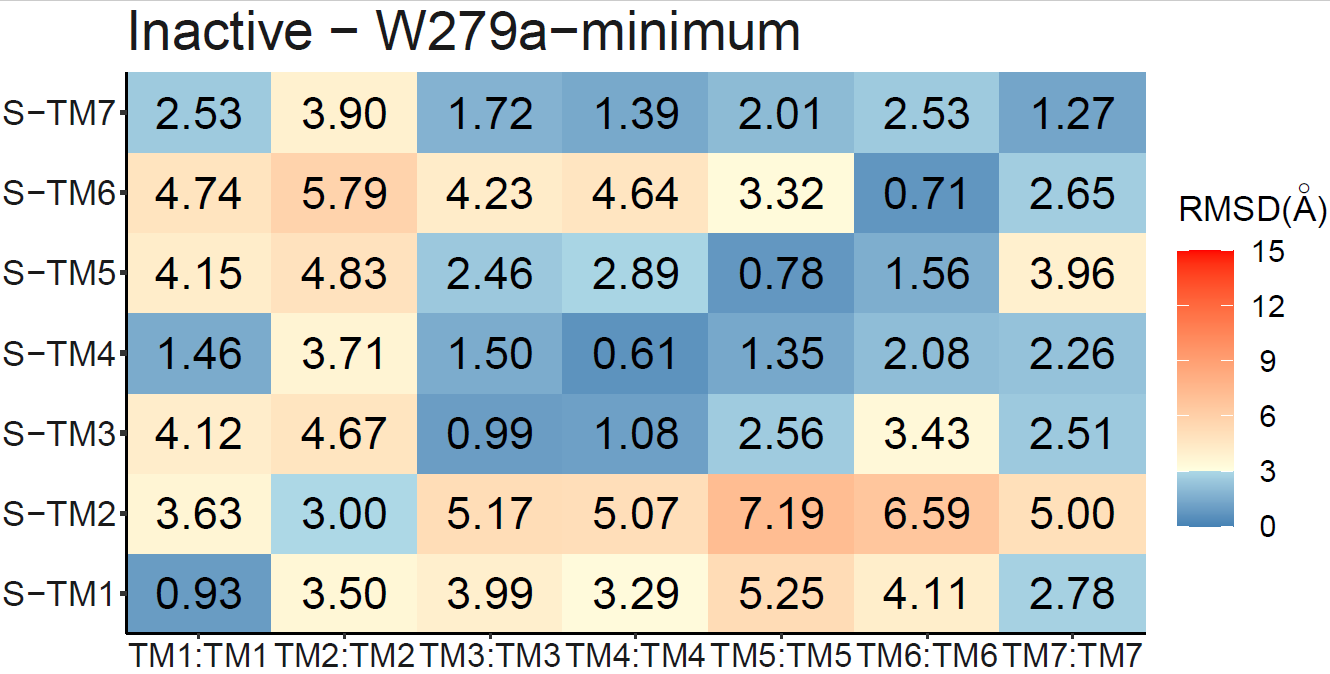

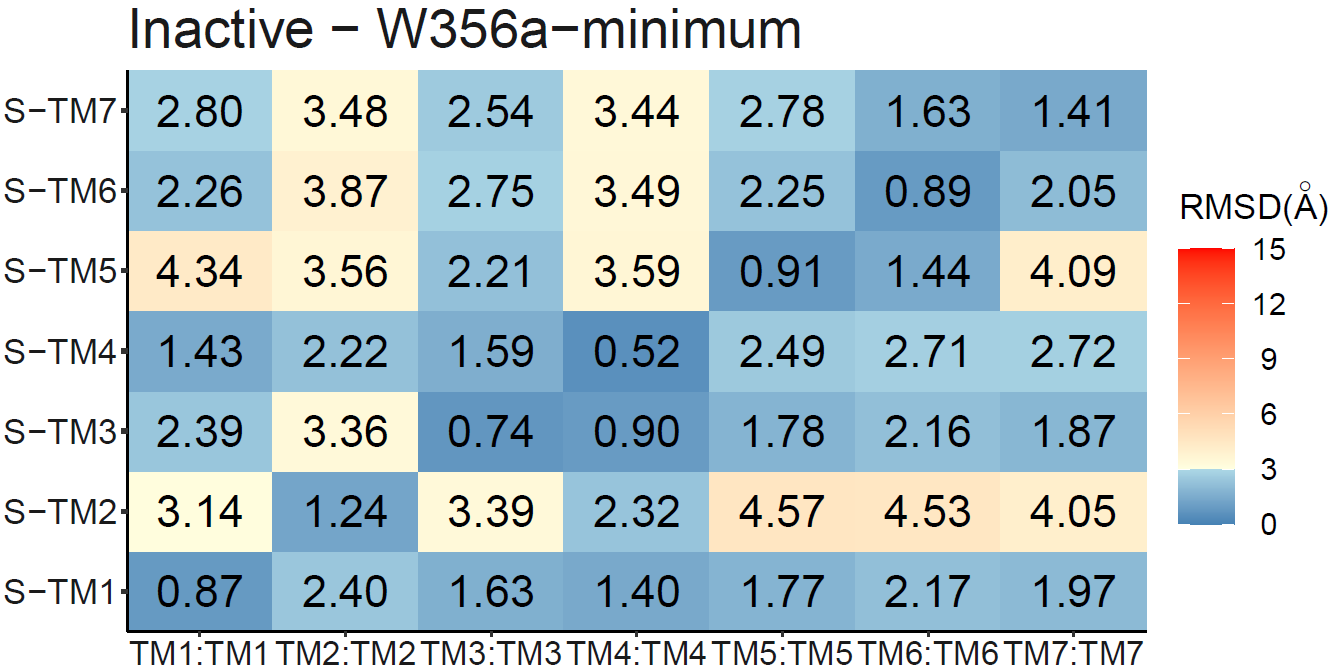

WT

W279A

F200A+W356A

W356A

F200A

F200A+W279A

W279A+W356A

F200A+W279A+W356A

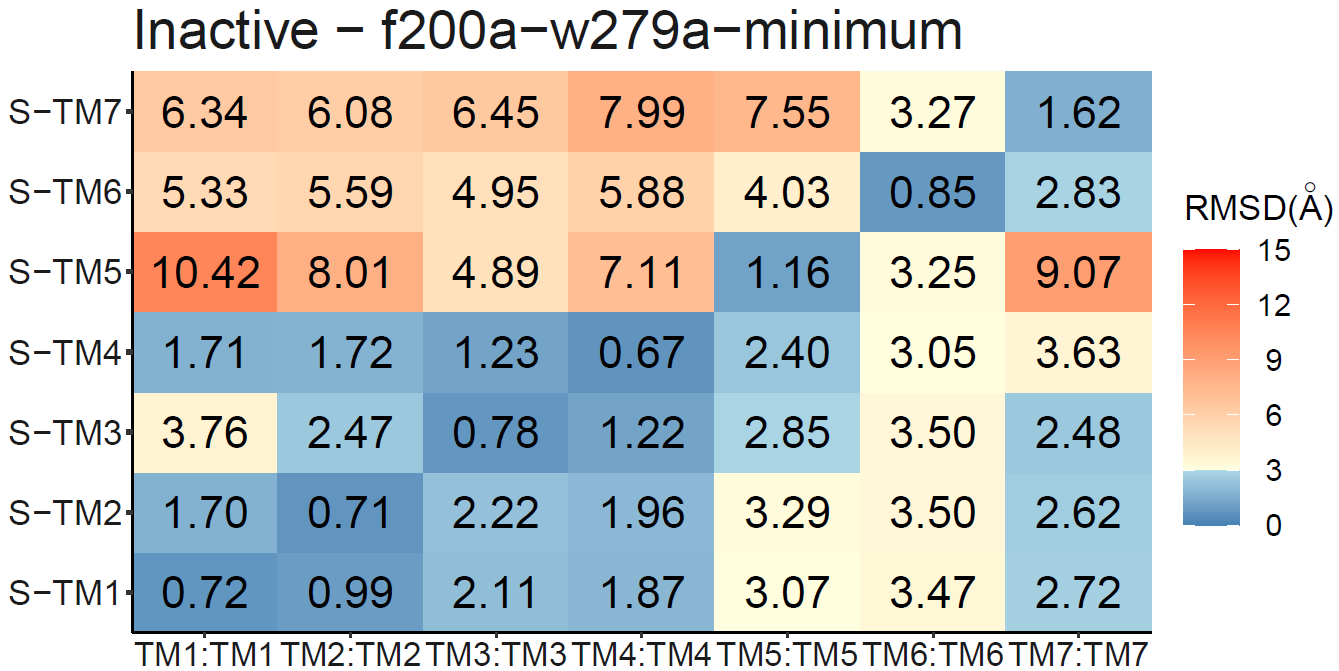

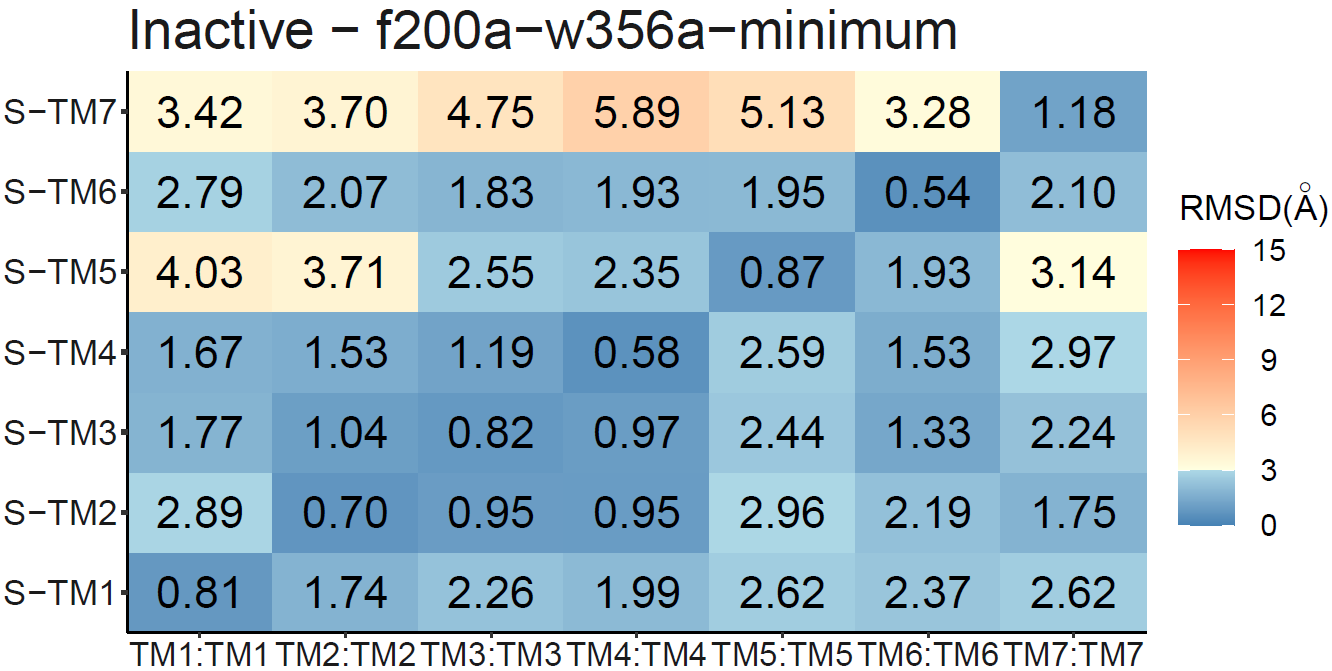

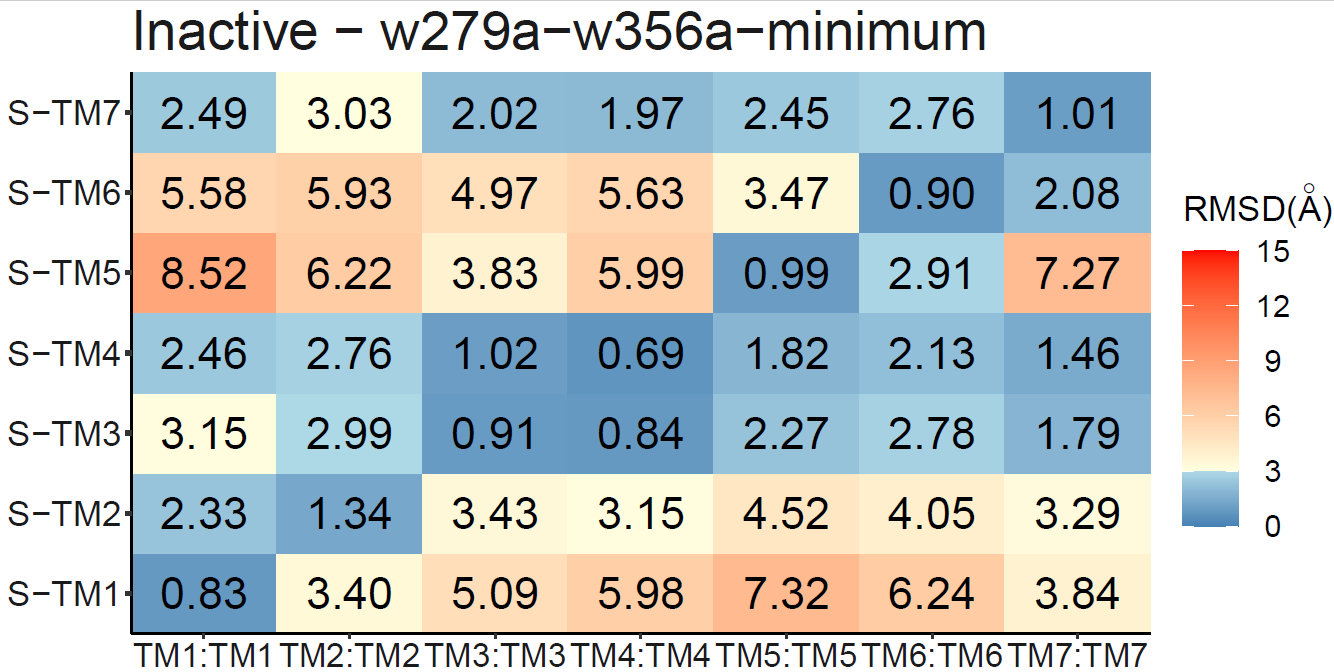

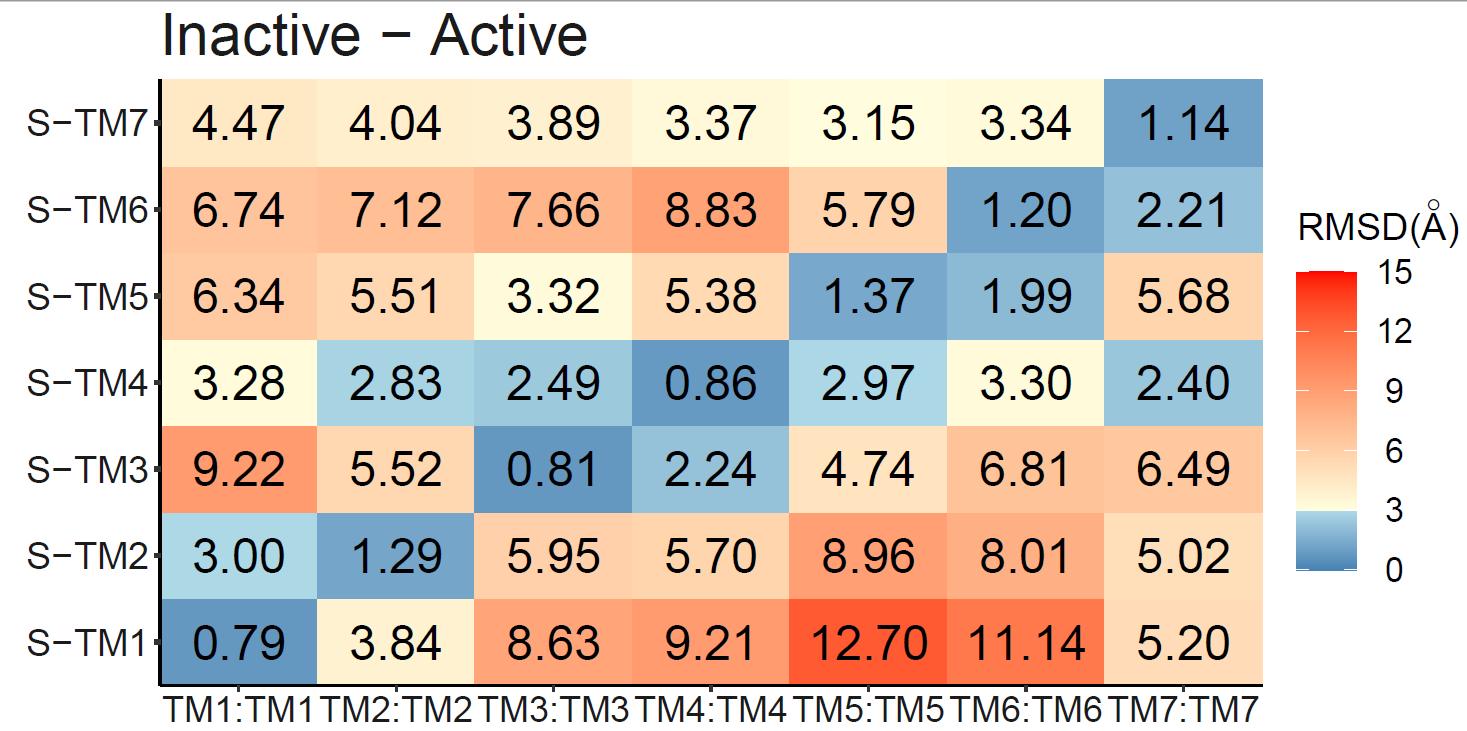

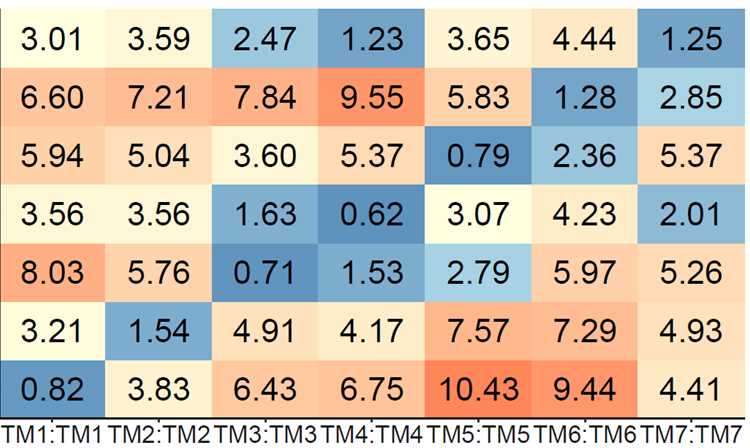

**Figure S3. 7×7 pairwise RMSD matrix of transmembrane (TM) helices.** The analysis reveals that the relative orientations of TM helices deviate from their initial configurations in all systems, with notable shifts observed in W356A, F200A+W356A, and triple (F200A+W279A+W356A) mutants. These conformations are closer to the inactive state in terms of TM helix arrangement. The consistent presence of W356 in all three inactivating mutants highlights its critical role in maintaining the active-state helical architecture of CB1. Crystal structures of the inactive and active states are depicted in gray and yellow, respectively.

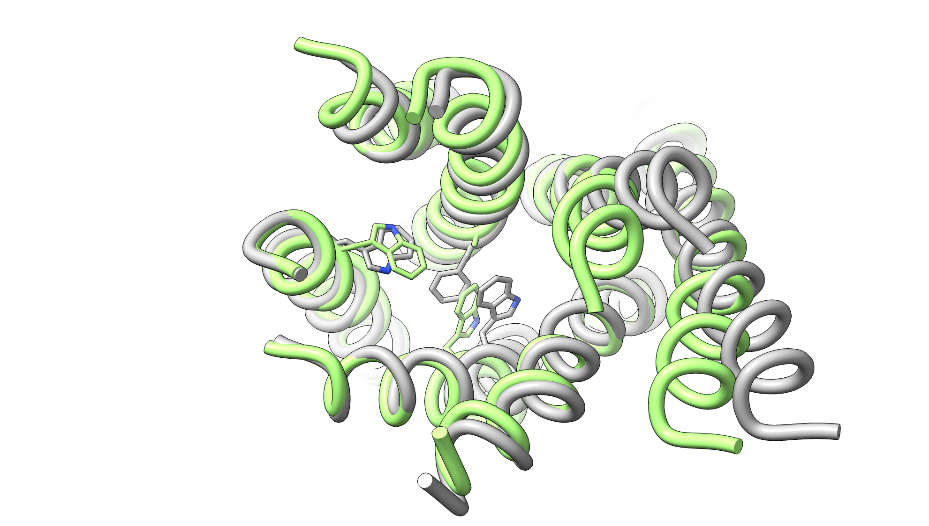

**W356**

**F200**

**W279**

F200A+W279A+W356A 1.639Å

F200A	RMSD=2.027Å

F200A+W356A	1.388Å

F200A+W279A	1.729Å

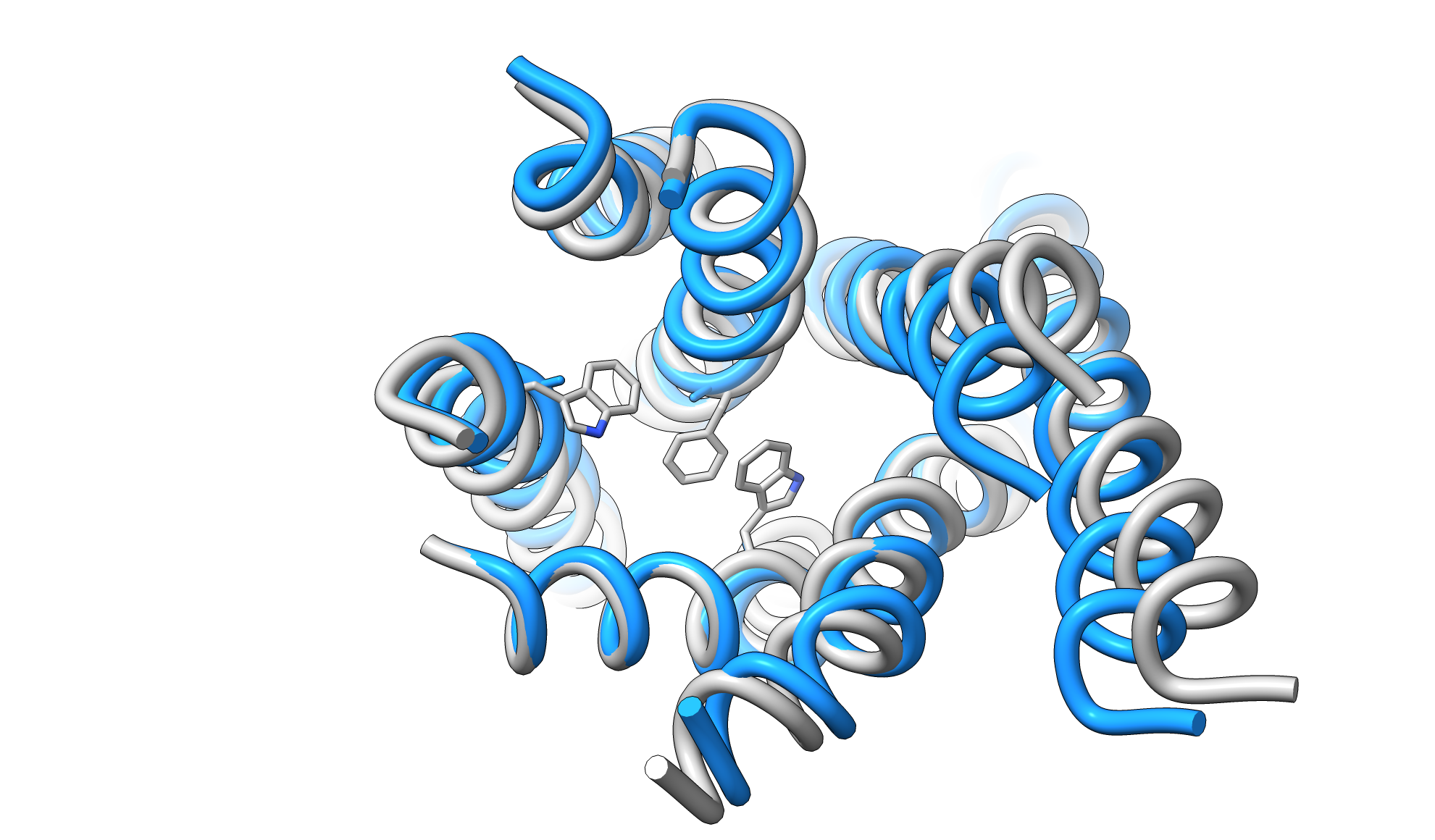

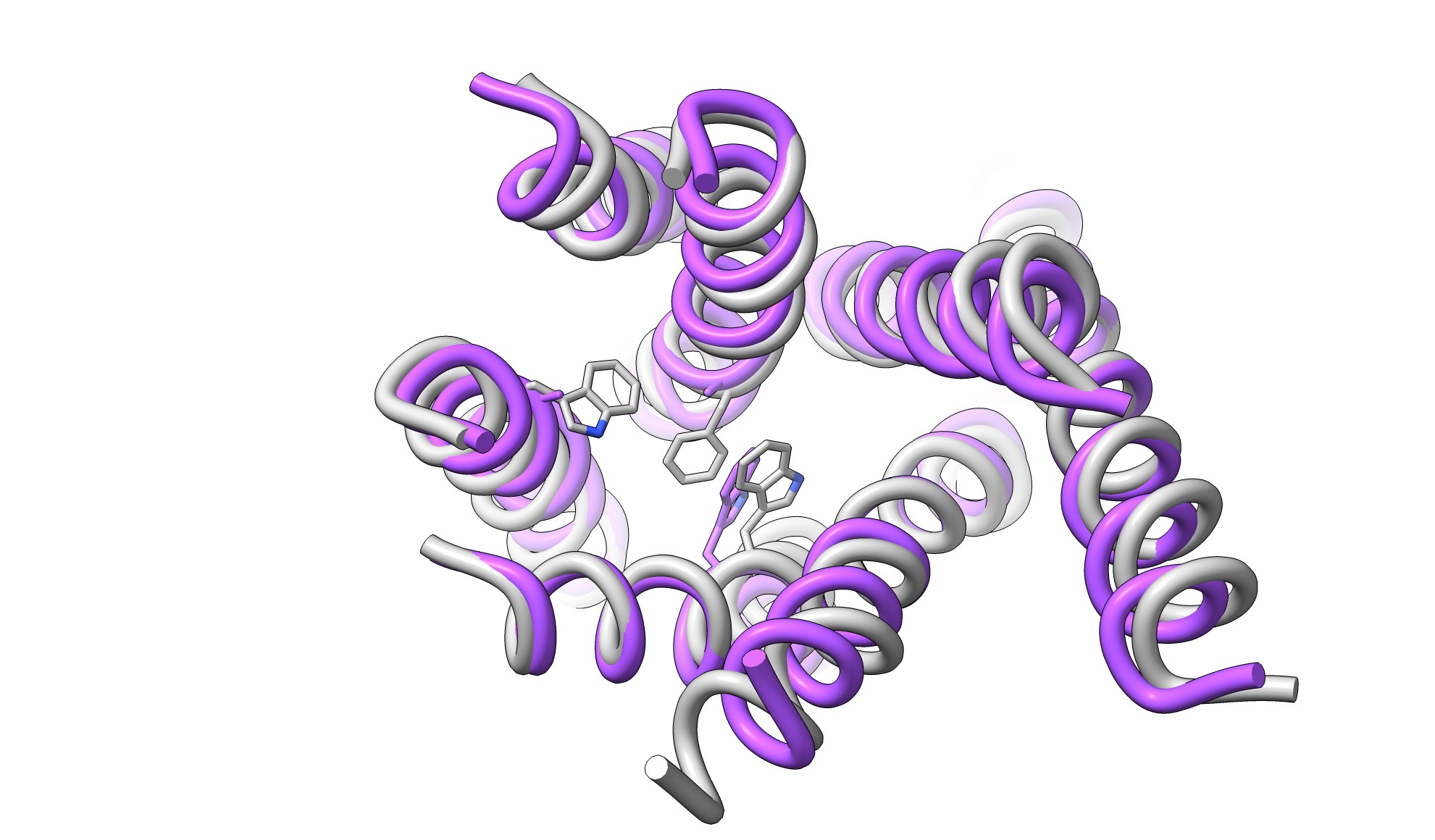

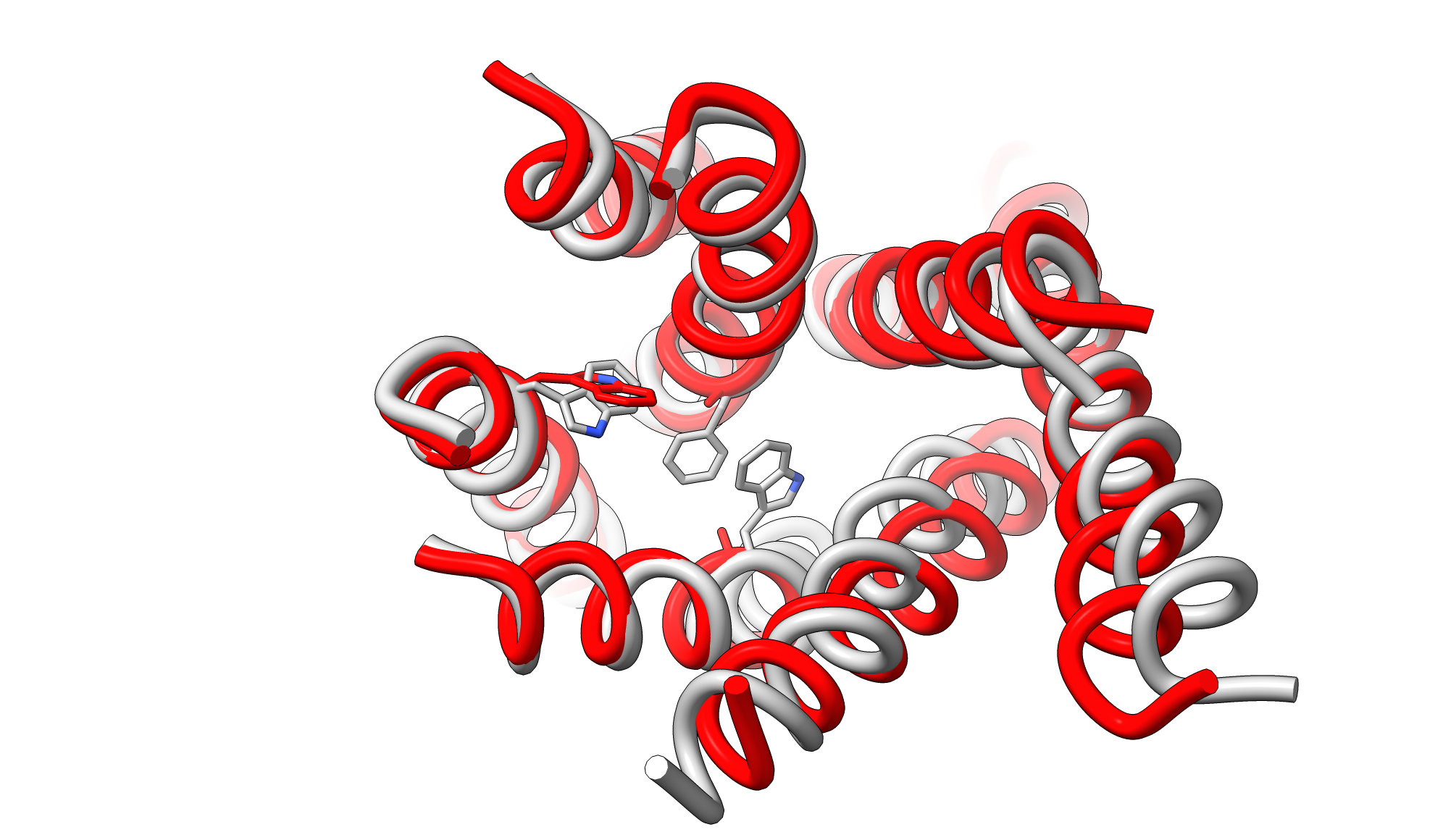

**W356**

**F200**

**W279**

**W356**

**F200**

**W279**

**W356**

**F200**

**W279**

**Figure S4. Aromatic microdomain residues conformations from simulation frames with the lowest RMSD to the inactive state.** Structures highlight conformational responses of the aromatic microdomain residues (F200^3.36^, W279^5.43^, W356^6.48^) to alanine mutations. In the F200A mutant, the A200 side chain adopts an orientation consistent with the inactive state, suggesting that this mutation facilitates toggle switch closure. In contrast, neither W279, W356, nor their respective alanine mutants adopt inactive-like conformations, indicating that these residues alone are insufficient to drive inactivation. These results underscore a dominant role for F200 in coordinating the transition toward the inactive state, whereas W279 and W356 may require cooperative interactions to follow.

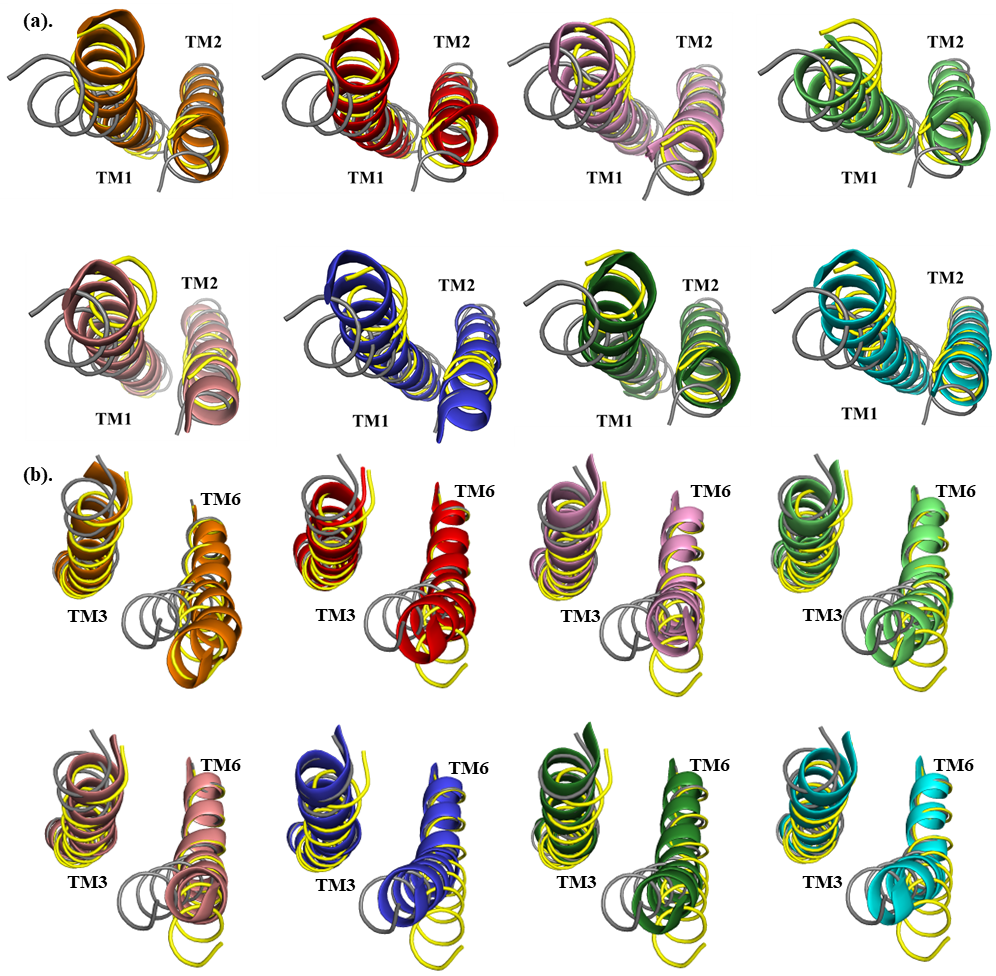
**Figure S5.** **Local structural comparisons of key transmembrane helices distinguishing active and inactive CB1 states.** **(a)** Extracellular view showing conformations of TM1 and TM2, the helices that display the most pronounced differences between active and inactive CB1 structures from mutant simulations most similar to the inactive conformation, overlaid with WT inactive (gray) and active (yellow) structures. **(b)** Intracellular view highlighting TM6, which exhibits the highest conformational flexibility and shift among all TMs across the simulations. Top row (left to right): wild-type active conformation, and single mutants F200A, W279A, and W356A. Bottom row (left to right): double mutants F200A+W279A, F200A+W356A, W279A+W356A, and the triple mutant F200A+W279A+W356A.

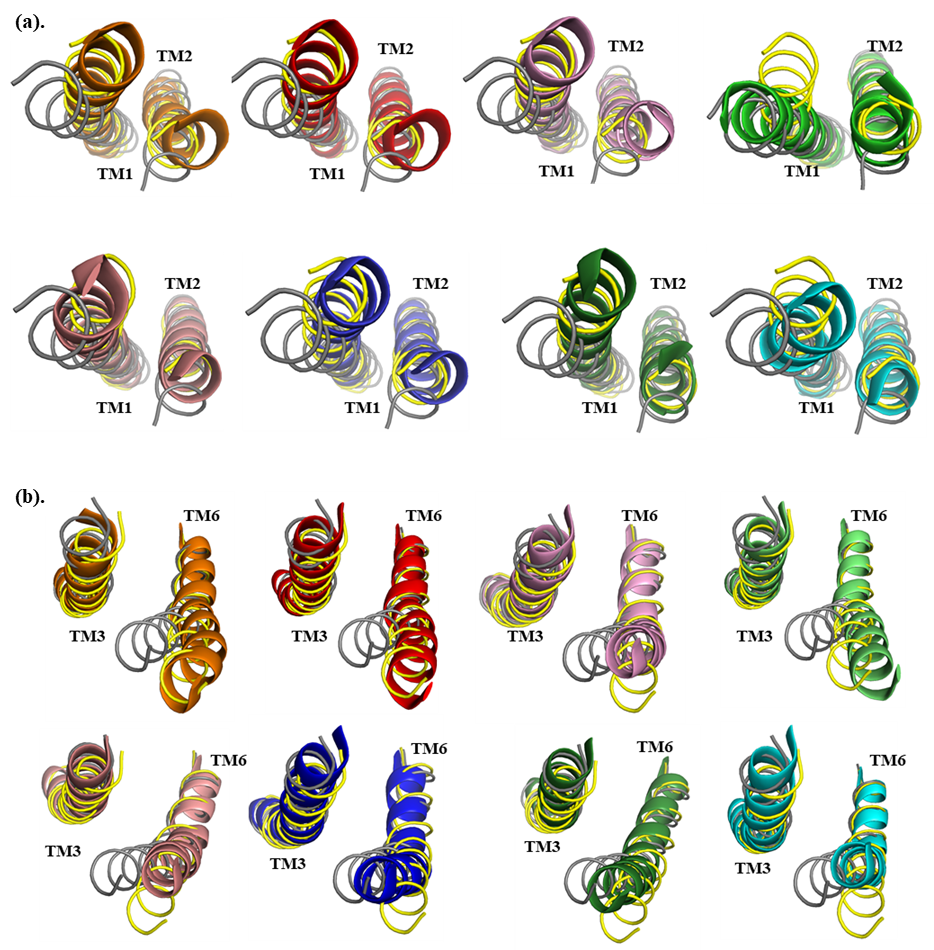

**Figure S6. Comparison of the most sampled conformations from simulations with the crystal structures of active and inactive CB1.**The dominant conformations sampled during simulations of each mutant are overlaid with the inactive (gray) and active (yellow) crystal structures. **(a)** The intracellular view highlights TM6 as the most conformationally flexible helix across systems. **(b)** The extracellular view reveals notable rearrangements in TM1 and TM2.From left to right, mutant structures are colored as follows: F200A (red), W279A (pink), W356A (lime), F200A+W279A (salmon), F200A+W356A (blue), W279A+W356A (green), and the triple mutant F200A+W279A+W356A (cyan).

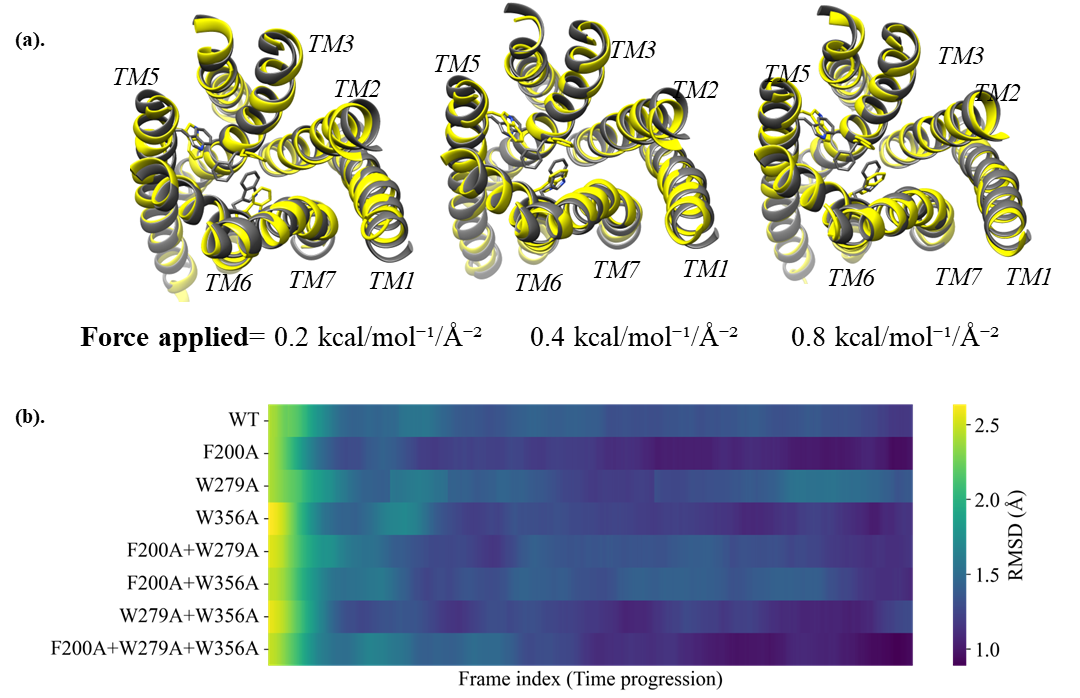

**Figure S7.** **Aromatic domain governs resistance to conformational transitions.** (**a)** RMSD values of WT CB1 relative to the inactive state following tMD simulations from the active-state conformation, with restraining forces applied to Cα atoms of TM helices at increasing magnitudes (0.2–0.8 kcal/mol⁻¹/Å⁻²). Higher forces progressively induced TM rearrangement, but F200^3.36^ retained its active-state rotamer in all cases, indicating decoupling of TM repacking and toggle switch transition. **(b)** RMSD values of WT and mutants CB1 after tMD simulations with minimal restraint (0.01 kcal/mol⁻¹/Å⁻²) applied to all TM atoms. Even under weak perturbation, all mutants adopted inactive-like conformations more readily than the WT, highlighting the role of these aromatic residues in stabilizing the active state.

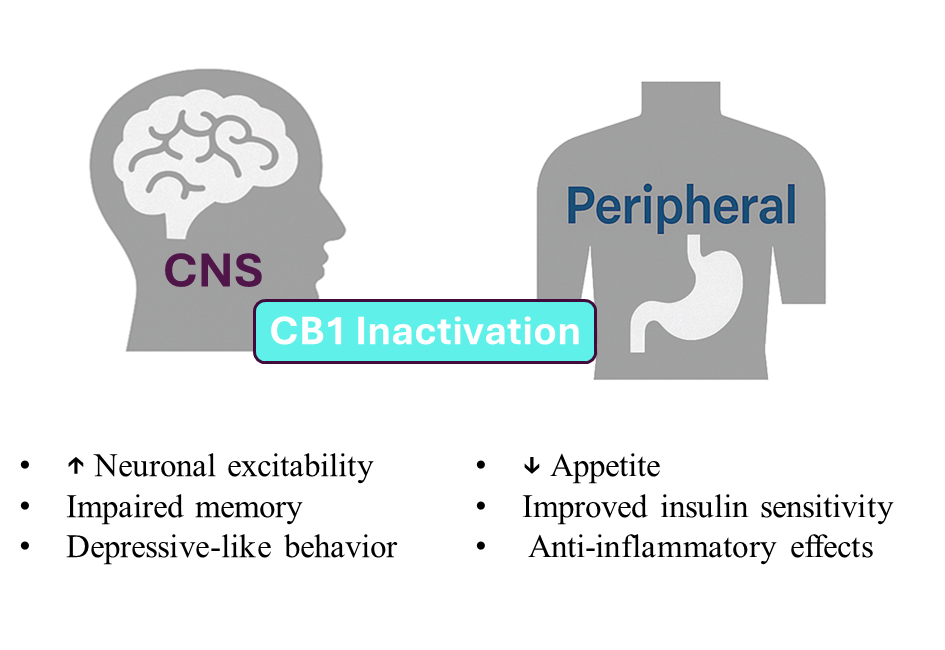

**Figure S8. Physiological consequences of CB1 inactivation bias across central and peripheral systems.** Schematic representation of how structural shifts in CB1 toward an inactive conformation via mutation or pharmacological modulation can influence distinct physiological processes. In the central nervous system (left), CB1 inactivation reduces endocannabinoid tone, which may result in heightened neuronal excitability, impaired memory consolidation, and increased vulnerability to anxiety or depressive behaviors. In peripheral tissues (right), CB1 inactivation is linked to reduced appetite, improved insulin sensitivity, and anti-inflammatory effects. These contrasting effects highlight the tissue-specific roles of CB1 and support the therapeutic potential of selectively modulating its conformational state.
